# Spatial and single-cell transcriptional landscape of human cerebellar development

**DOI:** 10.1101/2020.06.30.174391

**Authors:** Kimberly A. Aldinger, Zach Thomson, Parthiv Haldipur, Mei Deng, Andrew E. Timms, Matthew Hirano, Gabriel Santpere, Charles Roco, Alexander B. Rosenberg, Belen Lorente-Galdos, Forrest O. Gulden, Diana O’Day, Lynne M. Overman, Steven N. Lisgo, Paula Alexandre, Nenad Sestan, Dan Doherty, William B. Dobyns, Georg Seelig, Ian A. Glass, Kathleen J. Millen

## Abstract

Cerebellar development and function require precise regulation of molecular and cellular programs to coordinate motor functions and integrate network signals required for cognition and emotional regulation. However, molecular understanding of human cerebellar development is limited. Here, we combined spatially resolved and single-cell transcriptomics to systematically map the molecular, cellular, and spatial composition of early and mid-gestational human cerebellum. This enabled us to transcriptionally profile major cell types and examine the dynamics of gene expression within cell types and lineages across development. The resulting *‘Developmental Cell Atlas of the Human Cerebellum’* demonstrates that the molecular organization of the cerebellar anlage reflects cytoarchitecturally distinct regions and developmentally transient cell types that are insufficiently captured in bulk transcriptional profiles. By mapping disease genes onto cell types, we implicate the dysregulation of specific cerebellar cell types, especially Purkinje cells, in pediatric and adult neurological disorders. These data provide a critical resource for understanding human cerebellar development with implications for the cellular basis of cerebellar diseases.

## INTRODUCTION

The cerebellum is a highly interconnected brain structure that is essential for coordinating motor function and modulating cognition, emotional regulation, and language.^1^’^3^ Cerebellar development occurs over a broad time window that requires precise molecular regulation among multiple cell types.^4, 5^ Cerebellar dysfunction has been implicated in numerous diverse diseases. These include congenital structural abnormalities in children and cerebellar ataxias in adults that have specific cerebellar pathology, as well as disorders that impact multiple brain regions including the cerebellum, such as autism spectrum disorders (ASD) and Alzheimer’s disease.^1,6-8^ Though relatively well understood in mice,^9,10^ our molecular understanding of the complexity and dynamics of the human cerebellar development is very limited, with potentially human-specific expansion of progenitor zones.^11^

Cerebellar spatial organization results from timed cellular differentiation within distinct progenitor niches and coordinated cell migration during development.^11^’^13^ The cerebellar anlage emerges from the dorsal hindbrain and contains two primary proliferative niches: the ventricular zone (VZ) lining the fourth ventricle and the rhombic lip (RL), dorsal to the VZ, adjacent to the developing choroid plexus. The VZ, which gives rise to GABAergic neurons, is active during early embryonic stages and by 10 post conceptional weeks (PCW), nascent VZ-derived Purkinje cells dominate the cerebellar anlage **(Fig. 1a)**. After 10 PCW, the rhombic lip expands and generates glutamatergic neurons, including a multitude of granule cell progenitors, which migrate rostrally to constitute the external granule cell layer (EGL) on the dorsal surface. By mid-gestation, Purkinje cells reorganize to form a Purkinje cell layer (PCL) under the EGL, while granule cell progenitors within the EGL proliferate, differentiate, and migrate radially inwards, to form the internal granule cell layer located just below the Purkinje cell layer.

**Fig. 1.**
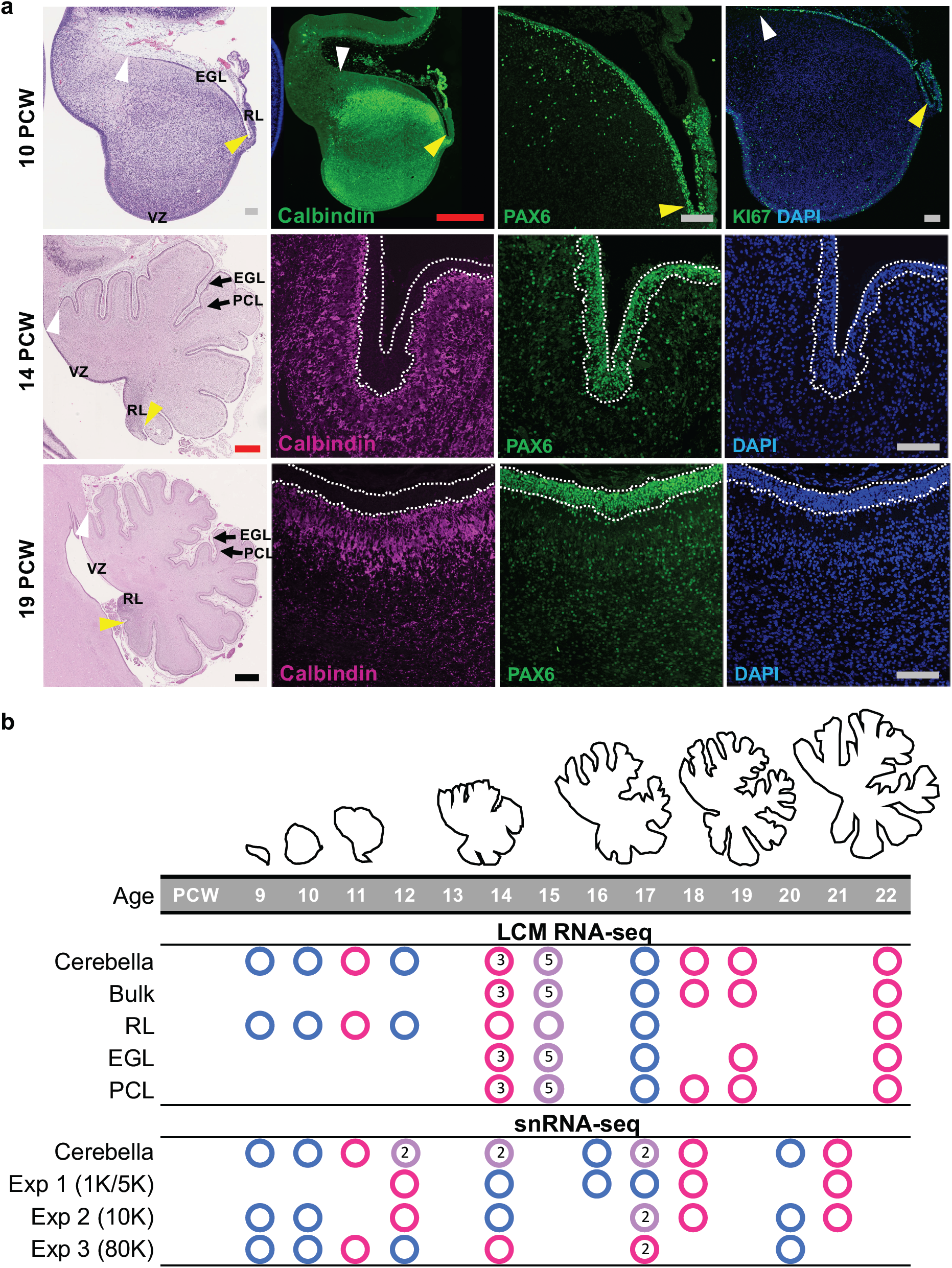
Overview of prenatal cerebellar development and the data generated in this study. **a**, Midsagittal sections of the human fetal cerebellum stained with hematoxylin and eosin (H&E) or markers for Purkinje cells (Calbindin) or rhombic lip (RL) and external granule cell layer (EGL; PAX6 and KI67). The ventricular zone (VZ), RL, EGL, and Purkinje cell layer (PCL) are shown. Arrowheads mark the anterior (yellow) and posterior (white) EGL across the dorsal surface of the cerebellar anlage. At 9 PCW, the cerebellar anlage is dominated by Purkinje cells, with a thin nascent EGL extending from the RL. By 19 PCW, Purkinje cells have migrated radially to establish a multicellular layer (PCL) beneath the EGL. Scale bars: 100 um (grey), 500 um (red), 1 mm (blue). *10 PCW specimen was used previously in Haldipur et al*., *2019*. **b**, The time span of fetal cerebellar development represented by line drawings of midsagittal sections of the cerebellum (to-scale) showing a dramatic change in size and foliation from 9 to 22 PCW. Below is the distribution of samples in this study. Each circle represents a cerebellum, color indicates sex [pink (female), blue (male), purple (at least 1 female and 1 male)], number indicates where >1 cerebellum was included.

The extensive growth and coordinated reorganization during cerebellar development is guided by molecular and cellular cues.^4,9,10^ However, cerebellum is not well represented in previous bulk and single-cell transcriptomic studies of the developing human brain.^14^’^18^ In the BrainSpan atlas of the developing human brain, cerebellum accounts for only 6% (35/607) of the total samples included **(Extended Data Fig. 1)**. In contrast, samples from neocortical regions account for 68% (415/607) of the samples included in this resource.^16^ Prenatal development of the cerebellum is even less well represented in BrainSpan, with the entirety of cerebellar development comprising just 5% (13/261) of the prenatal samples [8-37 postconceptional weeks (PCW)]. Further, the available data are predominantly derived from bulk transcriptomic analysis. Thus, the available human brain transcriptomic data are insufficient to capture the depth and breadth of the molecular repertoire present within the cerebellum, especially during early development, when developing Purkinje cells dominate the cerebellar anlage.

Here, we characterize the transcriptional and cellular landscape of the developing human cerebellum by using laser capture microdissection (LCM) to profile the transcriptomes of spatially defined progenitor and neuronal populations. We further employed single-cell combinatorial indexing to profile the transcriptomes of 70,000 cells across prenatal cerebellar development from 9-22 PCW. Finally, we compared our datasets to bulk RNA-sequencing expression profiles in the BrainSpan dataset. Our data establish a *‘Developmental Cell Atlas of the Human Cerebellum’* as a foundation for understanding developmental biology and to provide opportunities to explore disease impact to this brain structure.

## RESULTS

### Study design and data generation (Fig. 1)

We generated and analyzed transcriptomic data using direct and inferred methods for defining cell populations to characterize the transcriptional landscape of the prenatal human cerebellum. We used LCM to isolate cells within spatially demarcated progenitor and neuronal regions together with bulk RNA-seq (57 samples from 16 cerebella) and single-cell RNA-seq (69,174 cells/nuclei from 13 cerebella) from 29 postmortem cerebella obtained from clinically and histopathologically unremarkable donors of both sexes across fetal development **(Fig. 1b and Supplementary Tables 1-2)**.

To analyze the transcriptome of the primary progenitor and neuronal populations in the developing human cerebellum, we devised an experimental procedure to isolate cells occupying the RL, EGL and PCL **(Fig. 1a, Fig. 2a, and Extended Data Fig. 2a)**. Specifically, we dissected whole cerebella from fetal specimens with intact calvaria to ensure correct orientation of the cerebellum. We sectioned frozen cerebella in the sagittal plane through the cerebellar vermis, isolated RNA from one section (referred to from here on as ‘bulk’), and assessed RNA quality for each specimen [RNA integrity number (RIN), 7.7±0.95 (meanls.d.)] **(Supplementary Table 1)**. The EGL was easily identified histologically in sagittal sections as a cell-dense layer on the dorsal surface of the developing cerebellar anlage **(Fig. 1a and Supplementary Fig. 2a)**. To confirm the PCL, we performed immunohistochemistry (IHC) on an adjacent section using an anti-Calbindin antibody, a well-known Purkinje cell marker **(Fig. 1a and Supplementary Fig. 2a)**. Finally, we performed LCM to capture enriched neuronal populations from the EGL and PCL, isolated total RNA, and performed RNA-seq. We previously performed LCM and RNA-seq of the RL.^11^

**Fig. 2.**
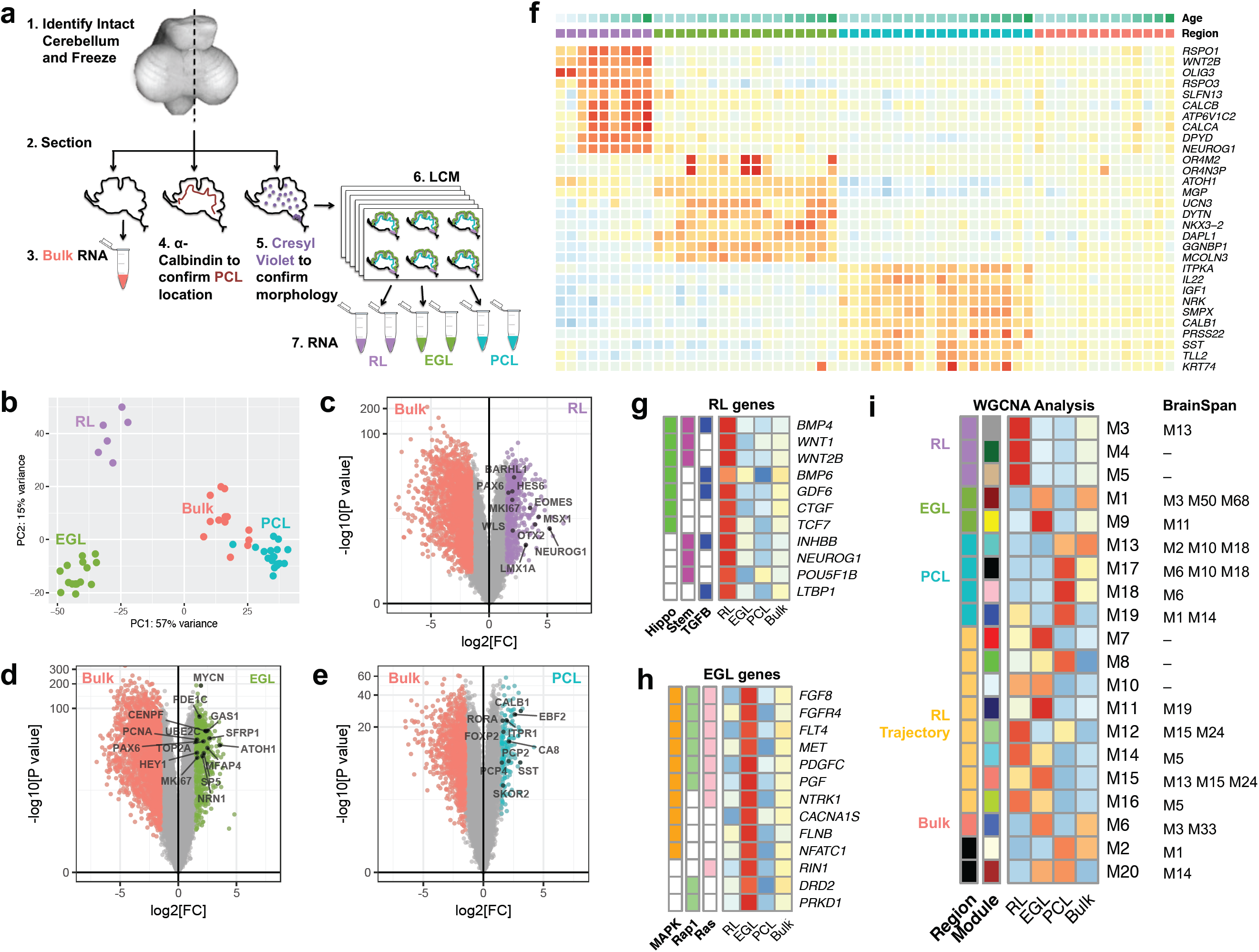
Spatial transcriptional analysis of the developing human cerebellum. **a**, Schematic illustrating LCM experimental workflow. **b**, Principal component analysis indicates that the largest source of variation among RNA-seq samples was spatial location, accounting for 57% of the variance, and verifying LCM successfully captured these regions. **c-e**, Volcano plots illustrating differential expression of genes for each spatial region versus bulk cerebellum. Colored dots represent genes with significant expression [FDR<0.05; Log2(FC)>1.5]. Selected canonical genes with significant expression are labeled. **f**, Heatmap of the top 10 genes with significant expression per spatial region (RL, EGL, PCL) are shown for each sample. Samples are ordered by region [RL (purple), EGL (green), PCL (turquoise), bulk (salmon)], then by ascending age (9 to 22 PCW). High expression is in red and low expression is in blue. **g**,**h**, Heatmaps of genes and pathways expressed in RL (g) and EGL (h) identified by gene ontology analysis. **i**, Heatmap of genes expressed in each WGCNA module enriched in **a**.

To prepare sequencing libraries, we used the Illumina TruSeq RNA Access Library Prep Kit because it requires low total RNA input, yet maintains high sensitivity for transcripts expressed at low levels. Paired-end Illumina high-quality sequencing was performed on 57 samples including bulk cerebellum (N=13), EGL (N=17), PCL (N=18), and RL (N=9) from 16 mid-gestation (9-22 PCW) fetal specimens **(Fig. 1b and Supplementary Table 1)**. We validated the enrichment of our LCM-isolation by comparing gene expression of established RL (*LMX1A, BARHL1)*, EGL (*ATOH1, PAX6)*, and PCL *[CALB1, SKOR2)* markers in the RNA-seq dataset from LCM-isolated samples and bulk-isolated cerebellum **(Extended Data Fig. 2b)**. The expression of these six neuron-specific markers confirmed the specificity of our enrichment, with the highest expression detected in the appropriate samples.

The spatially defined analyses were complemented by single nucleus RNA-seq (snRNA-seq) data generated across three experiments using 26 samples from an independent set of 13 cerebella ranging in age from 9 to 21 PCW **(Fig. 1b and Supplementary Table 2)**. We used split-pool ligation-based transcriptome sequencing (SPLiT-seq), a multi-step barcoding strategy combined with RNA-seq, to simultaneously interrogate thousands of cells/nuclei among multiplexed samples.^19^ Single-cell (8 samples) and single-nucleus (12 samples) level transcriptomic data were generated in technical replicates for 10 cerebella across two experiments; data for the remaining 3 cerebella were generated in a single experiment.

### Transcriptional analysis of spatially defined neural zones (Fig. 2)

To characterize the global transcriptional landscape in three spatially demarcated regions, we applied principal component analysis (PCA) to the expression profiles from LCM-isolated EGL, PCL, and RL samples and from bulk cerebellum. PCA showed that the samples clustered according to neuronal region, with PC1 distinguishing RL and EGL from PCL and bulk cerebellum samples and PC2 distinguishing RL from EGL **(Fig. 2b)**. To identify genes that are spatially regulated, we evaluated differential gene expression (DGE) between each LCM-isolated zone and bulk cerebellum using a modest threshold [Benjamini-Hochberg adjusted P value <0.05 and log2-transformed fold change >1.5] and including library prep batch, sex, and age as covariates **(Fig. 2b-d)**. There were 1,086 genes with significant DGE (1.5-fold, p-adj<0.05) between the neuronal zones: 620 genes showed increased expression in RL, 599 genes in EGL, and 163 genes in PCL compared to bulk cerebellum **(Fig. 2e and Supplementary Table 3)**. The RL genes were enriched for cell cycle (hsa04110, FDR = 6.66×10^−18^) and p53 signaling pathways (hsa04115, FDR = 5.44xl0’^−6^), as were the EGL genes (hsa04110, FDR = 6.71xl0’^−15^ and hsa04115, FDR = 0. 0006) **(Supplementary Table 4)**. The PCL genes showed no pathway enrichment. Subsets of genes were specifically expressed in each captured region **(Extended Data Fig. 3):** 181 RL-specific genes, 175 EGL-specific genes, and 136 PCL-specific genes. The RL genes were enriched in Hippo signaling, stem cell pluripotency regulation, and TGFb signaling **(Fig. 2f)**, with expression increasing across mid-gestation **(Fig. 2g)**. EGL genes were enriched in MAPK, Ras and Rap1 signaling **(Fig. 2h,i)**. No pathway enrichment was detected among PCL genes.

**Fig. 3.**
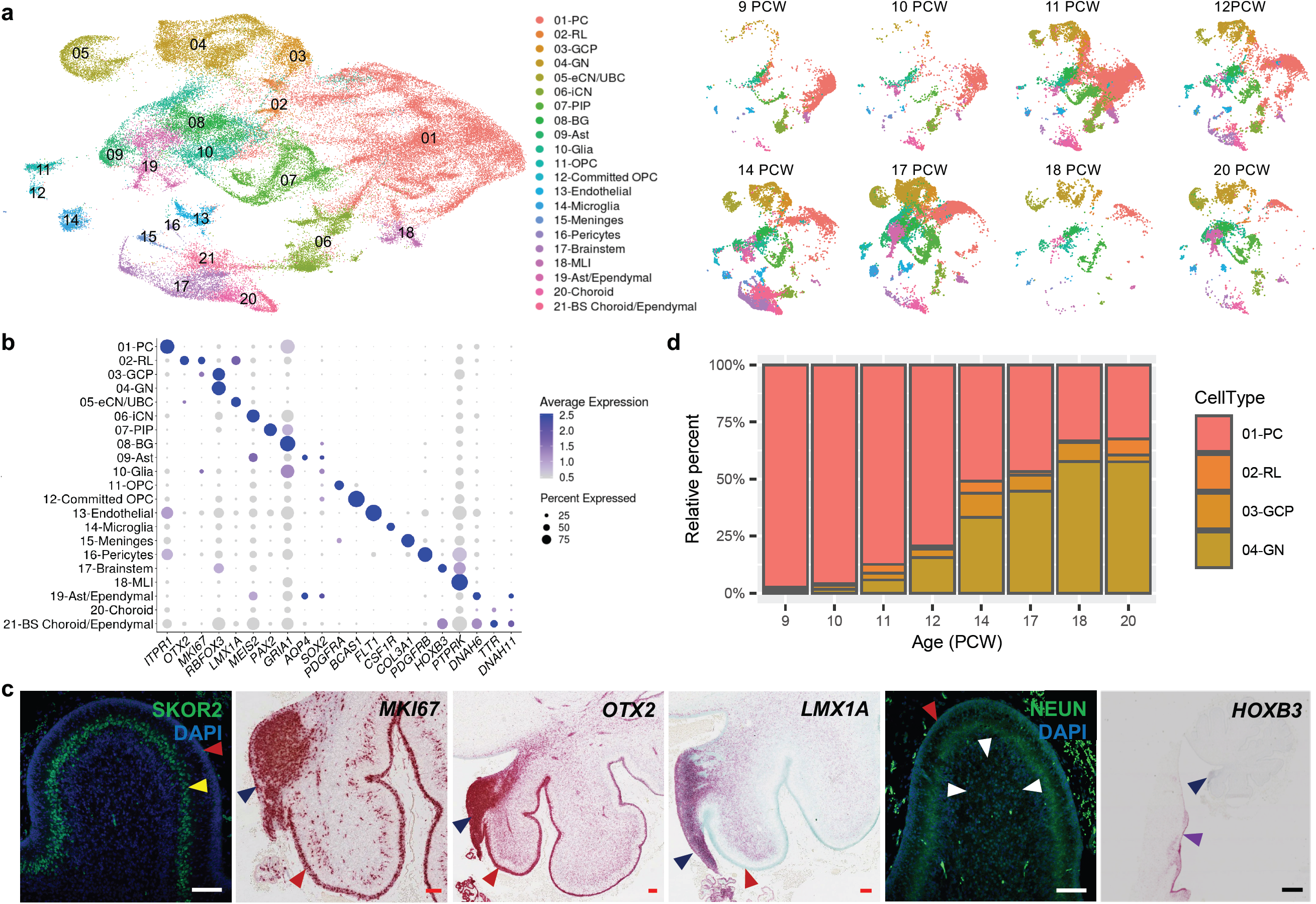
Identifying the major cell types of the developing human cerebellum. **a**, UMAP visualization of 67,174 human cerebellar nuclei colored by cluster identity from Louvain clustering and annotated on the basis of marker genes. The same UMAP is plotted at right, showing only nuclei from each age (nuclei numbers from left to right: n = 5,003 for 9 PCW; 2,329 for 10 PCW; 20,364 for 11 PCW; 7,119 for 12 PCW; 11,213 for 14 PCW; 15,556 for 17 PCW; 1,617 for 18 PCW; 5,177 for 20 PCW). **b**, Dot plot showing the expression of one selected marker gene per cell type. The size of the dot represents the percentage of cells within a cell type in which that marker was detected and its color represents the average expression level. **c**, Midsagittal sections of the human fetal cerebellum at 18 PCW stained with selected marker genes for Purkinje cells (SKOR2), proliferation (MKI67), Rhombic lip *(OTX2* and *LMX1A)*, granule neurons (NEUN) and brainstem *(HOXB3)*. The EGL, PCL, internal granule cell layer, RL and brainstem are indicated by red, yellow, white, blue and purple arrowheads, respectively. Sections are counterstained using DAPI for immunohistochemistry (SKOR2, NEUN) or Fast Green for *in situ* hybridization *(MKI67, OTX2, LMX1A, HOXB3)*. Scale bar = 100 um and 1 mm *(HOXB3). LMX1A was used previously in Haldipur et al*., *2019*. **d**, Percentage of the four major cell types from each age sampled. Bar colors represent Purkinje cells (PC), rhombic lip (RL), granule cell precursors (GCP), or granule neurons (GN).

To identify cellular components of the spatial cerebellar transcriptome, we performed weighted gene co-expression network analysis (WGCNA)^20^ on all 57 LCM samples and identified 21 modules of coexpressed genes **(Extended Data Fig. 4 and SupplementaryTables 5-6)**. We curated 21 gene coexpression modules according to spatial relationships between enriched regions and shared gene expression between regions within the RL lineage. We found 9 modules that showed expression differences among cerebellar regions (spatial), 8 modules that showed expression differences in both RL and EGL (RL lineage), one module enriched in bulk cerebellum, and three modules that did not show dynamic expression among the regions captured.

**Fig. 4.**
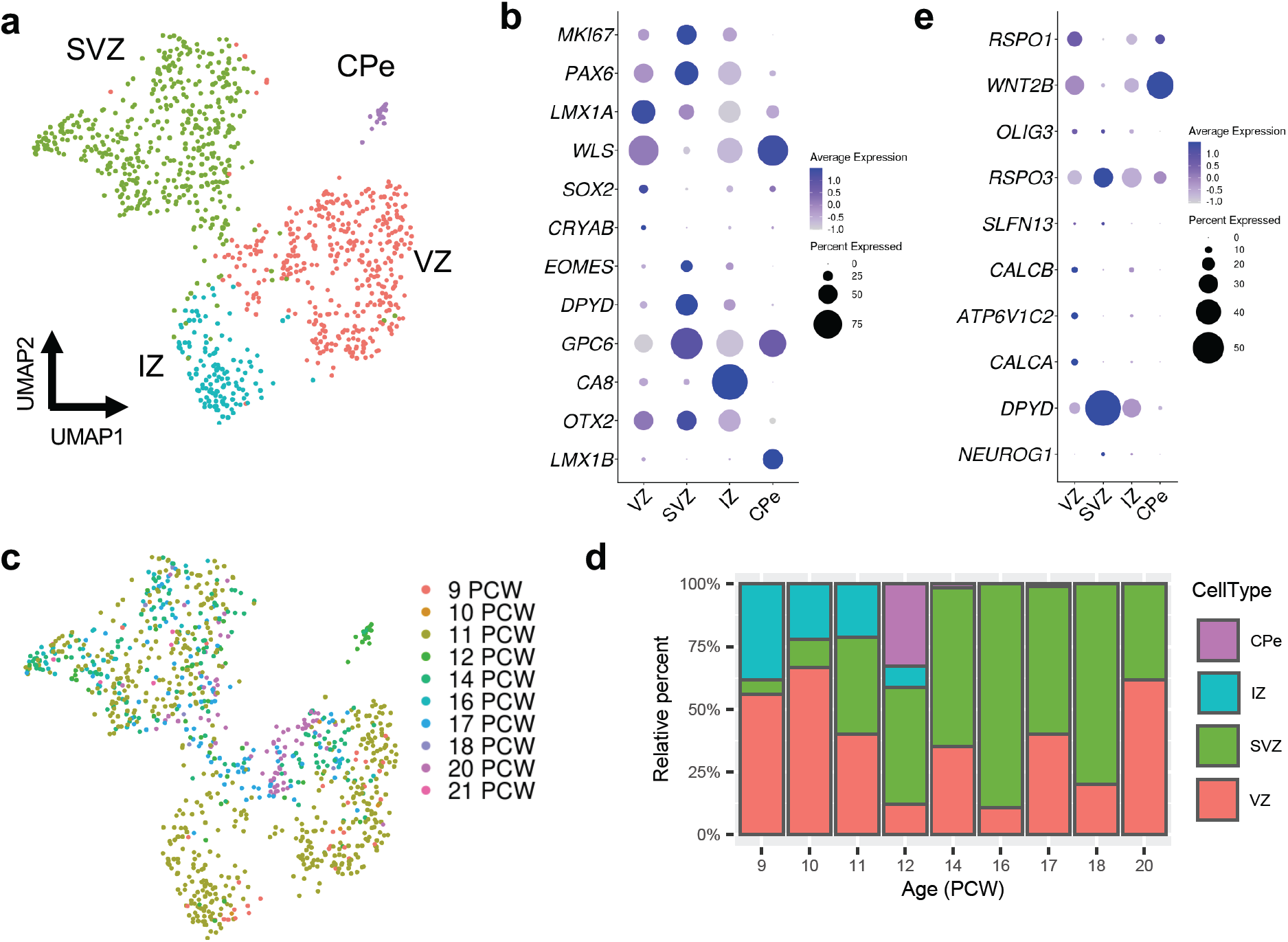
Analysis of RL compartments at single-cell resolution. **a**, UMAP visualization and marker-based annotation of the RL subclusters (n = 1,018 nuclei; 466 for SVZ; 390 for VZ; 135 for IZ; 21 for CPe). CPe, choroid plexus epithelium; IZ, intermediate zone; SVZ, subventricular zone; VZ ventricular zone, **b**, Dot plot showing the expression of selected marker genes in subclusters, **c**, The same UMAP as in **a** with nuclei colored by sample age (n = 34 for 9 PCW; 9 for 10 PCW, 535 for 11 PCW; 58 for 12 PCW; 137 for 14 PCW; 56 for 16 PCW; 97 for 17 PCW, 5 for 18 PCW; 81 for 20 PCW; 6 for 21 PCW). **d**, Percentage of the RL subclusters by sample age. **e**, Dot plot showing the expression of the top 10 most differentially expressed genes from the spatial transcriptional analysis of the RL.

We compared our 21 gene coexpression modules to the 73 modules generated in the most recent BrainSpan analysis of human neurodevelopment^16^ and found 26 of the 73 BrainSpan modules were correlated with the modules derived from our data **(Fig. 2j and Extended Data Fig. 5)**. We found that genes in 14 BrainSpan modules were enriched among genes with spatial expression in prenatal cerebellum, 8 were enriched with RL lineage, 2 were enriched bulk cerebellum, and 2 were correlated with modules that were not dynamic in the prenatal cerebellum. Overall, the majority of BrainSpan modules with genes that were enriched in our prenatal cerebellum modules were highly expressed prenatally in all brain regions and contained multiple neural and non-neural cell types **(Supplementary Table 3)**. Among the 15 BrainSpan cerebellar modules, only one (Mil) was shared with our data (M9). In BrainSpan, Mil is highly expressed in postnatal cerebellum, and includes granule cell markers, including *PAX6* and *GABRA6*. This result was expected since the BrainSpan dataset contains few cerebellum samples overall and those that are present are dominated by postnatal ages when granule cells vastly overwhelm all other cell types in the cerebellum,^21^ while our data are exclusively prenatal. In our data, *PAX6* is found in M14, which is highly expressed in both RL and EGL and enriched in processes regulating DNA **(Supplementary Table 6)**.

**Fig. 5.**
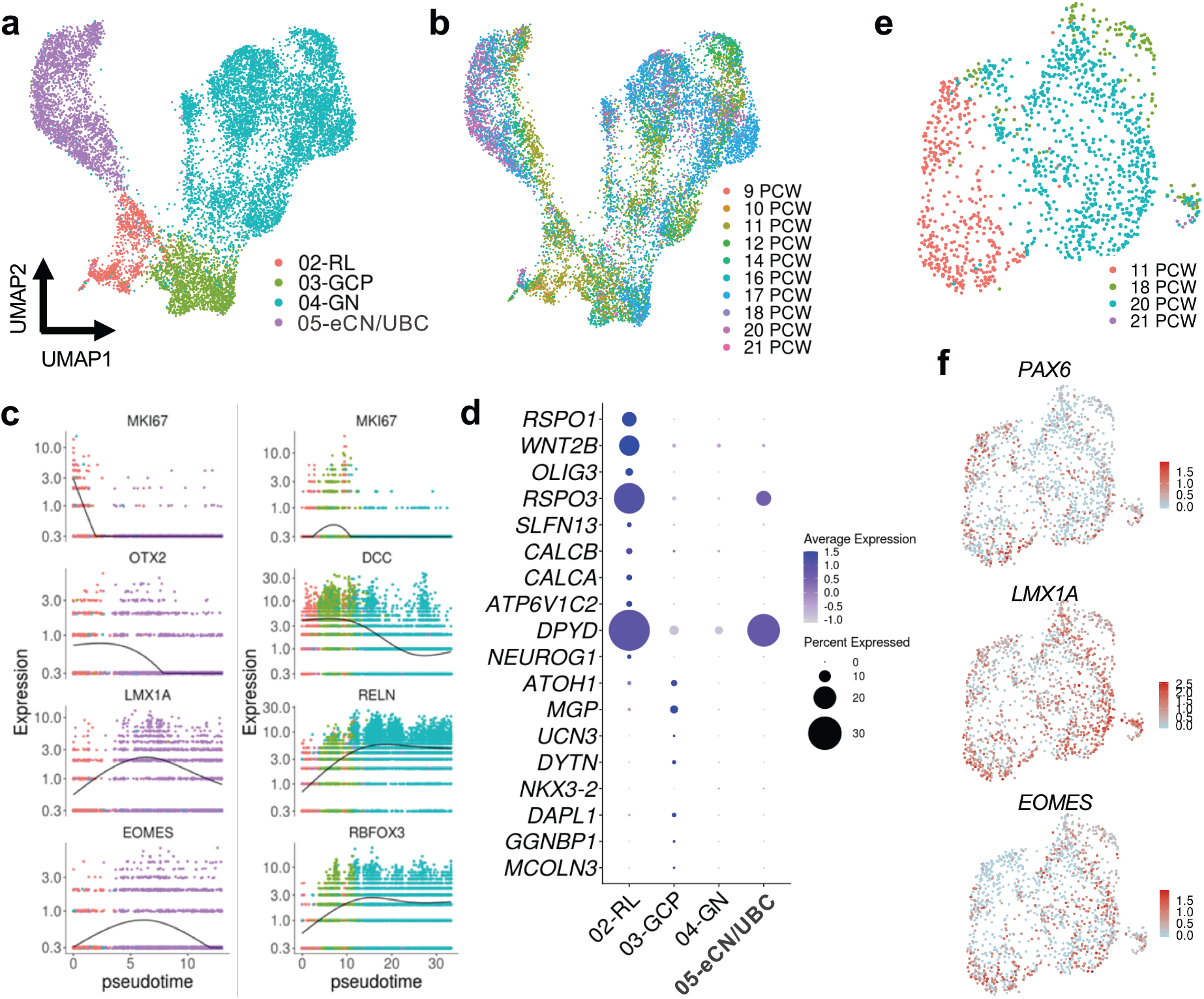
Characterization of the RL trajectory. **a**, UMAP visualization and marker-based annotation of cell types that originate from the RL (n = 12,243; 1,018 for RL; 1,659 for GCP; 6,727 for GN; 2,839 for eCN/UBC). eCN/UBC, excitatory cerebellar nuclei/unipolar brush cells; GCP, granule cell progenitors; GN, granule neurons; RL, rhombic lip. **b**, The same UMAP as in **a** with nuclei colored by sample age. **c**, Kinetics plot showing the relative expression of RL trajectory marker genes across developmental pseudotime. Dots are colored according to cell types as in **a. d**, Dot plot showing the expression of the top 10 most differentially expressed genes from the spatial transcriptional analysis of RL and EGL. **e**, UMAP visualization of the eCN/UBC cluster including 11, 18, 20, 21 PCW samples. Nuclei are colored by sample age. **f**, The same UMAP as in **e** with nuclei colored by expression level for the indicated gene.

### Cell types in the developing human cerebellum (Fig. 3)

We performed single-nucleus RNA-seq to define cell types and assemble cell-type transcriptomes in the developing human cerebellum from 9 to 21 PCW **(Supplementary Table 2)**. Using SPLiT-seq,^19^ we successfully sequenced a total of 92,314 nuclei with a median transcript capture of 1,214 unique molecular identifiers (UMIs) per profile **(Supplementary Table 7)**. We removed outlier cells with too few (<200) or too many (dataset specific) genes detected. We used DoubletFinder^22^ to detect 5% likely doublet cells that we subsequently discarded. The 69,174 nuclei analyzed had an average of 3,626 transcripts/UMIs per nucleus from 1,332 genes. A total of four datasets from three experiments **(Fig. 1b)**, including two smaller datasets generated previously,^23^ and two additional datasets generated through independent experiments herein, were merged for the analysis. Each dataset was filtered independently, then we used Seurat v3^24^ to integrate all four datasets. We applied Louvain clustering and UMAP visualization to all cells in the integrated dataset **(Fig. 3a)**. Cells from replicate samples processed in separate experiments were similarly distributed while cells from different developmental stages were not **(Extended Data Fig. 6)**. Using known cell type marker genes to manually annotate clusters, we validated the expression of selected marker genes using immunohistochemistry *or in situ* hybridization, and identified 21 distinct cell types **(Fig. 3b**,**c and Supplementary Table 8)**.

**Fig. 6.**
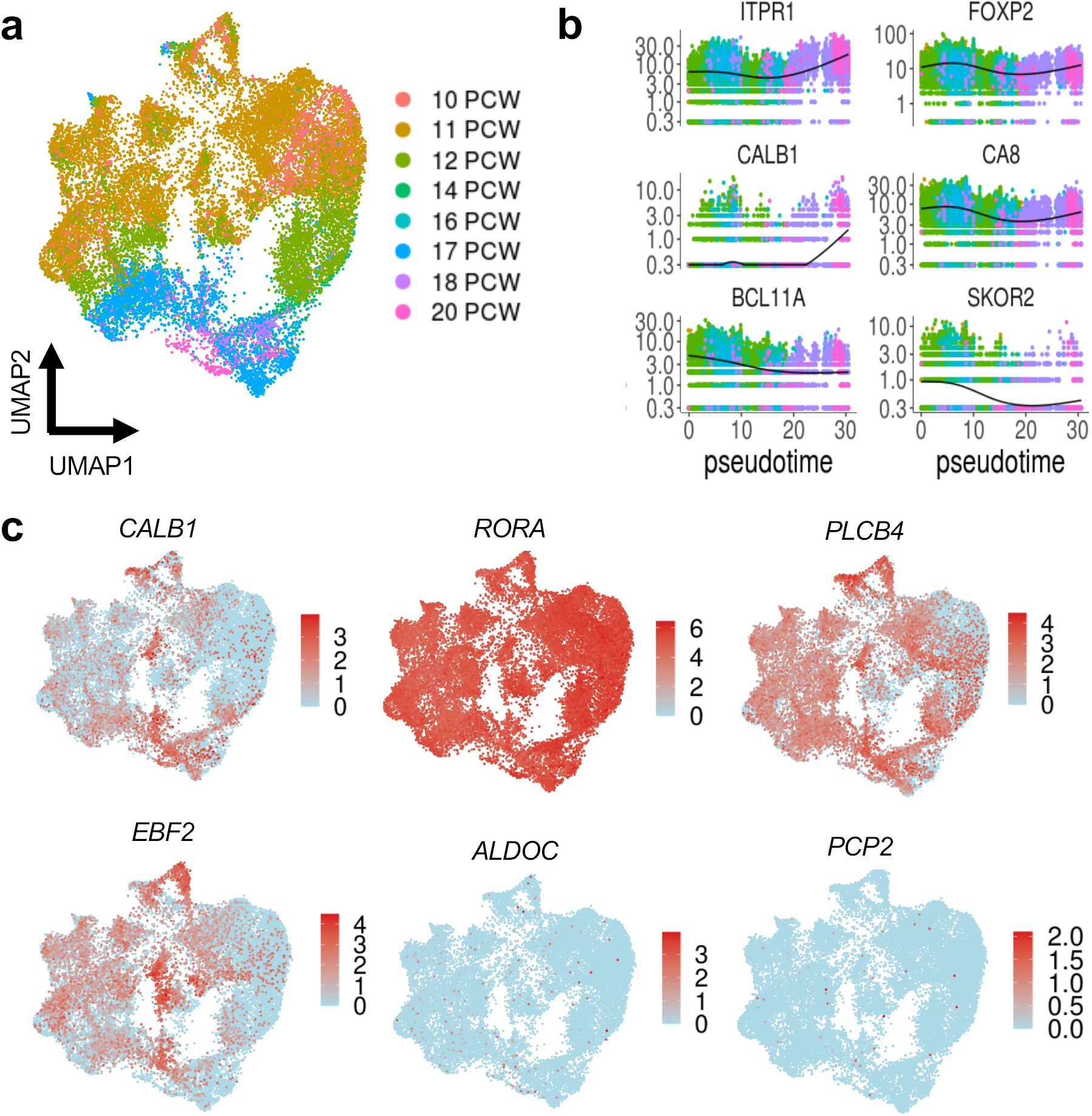
Purkinje cells. **a**, UMAP visualization of the PC cluster (25,724 nuclei). Nuclei are colored by sample age. **b**, Kinetics plot showing the relative expression of PC marker genes across developmental pseudotime. Dots are colored by sample age as in **a. c**, The same UMAP as in **a** with nuclei colored by expression level for the indicated gene.

Across the 21 major cell types, 4,443 genes (FDR < 0.05) were differentially expressed **(Supplementary Table 9)**. We identified 239 cell-type specific marker genes **(average LogFC>1**.**5; Extended Data Fig. 7)**, many of which were previously characterized as markers of the respective cell types. For example, we detected *CA8, ITPR1, DAB1, and RORA* in Purkinje cells, *5LIT2* in the rhombic lip, and *RELN* and *RBFOX3* in granule neurons.

**Fig. 7.**
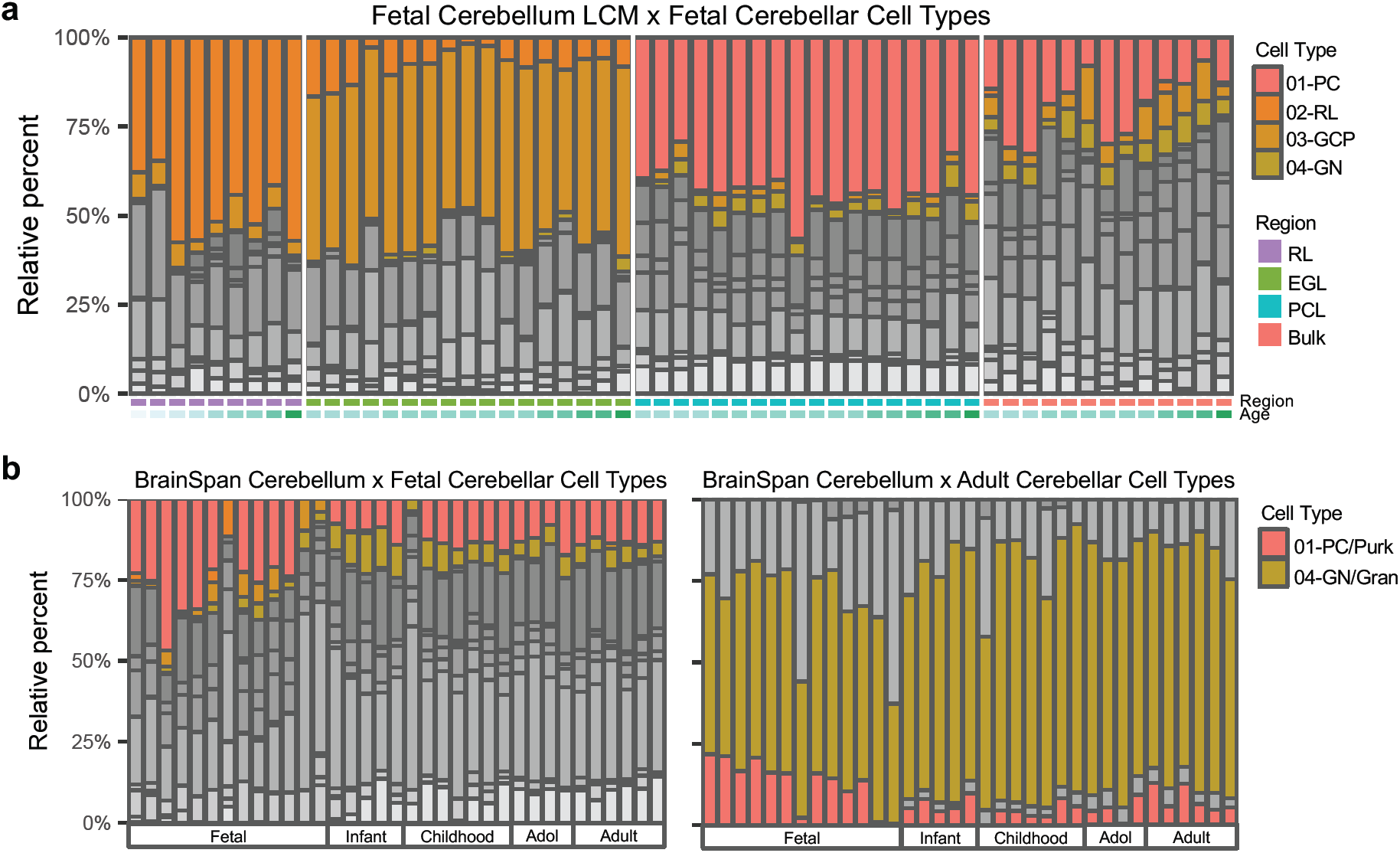
Inferred cell type composition of bulk cerebellum RNA-seq. **a**, Percentage of the four major fetal cerebellar cell types in each sample of the spatial transcriptional analysis (n=57). Bar colors represent Purkinje cells (PC), rhombic lip (RL), granule cell precursors (GCP), or granule neurons (GN). Additional cell types are in grayscale. Samples are ordered by region [RL (purple), EGL (green), PCL (turquoise), bulk (salmon)], then by ascending age (9 to 22 PCW). **b**, Percentage of the four major fetal cerebellar cell types (left) or the two major postnatal cerebellar cell types (right) in each cerebellar sample from BrainSpan (n=35). Bar colors represent fetal (01-PC) or adult (Purk) Purkinje cells and fetal (04-GN) or adult (Gran) granule neurons. Additional cell types are in grayscale. Samples are ordered by ascending age (8 PCW to 40 years). Developmental stage of the samples is shown below the bar plots.

The 21 cell types are represented by a median of 1,659 nuclei, the largest containing 25,724 nuclei (Purkinje cells) and the smallest containing only 189 nuclei (pericytes). The four major cell types (Purkinje cells, rhombic lip, granule cell progenitors, granule neurons) in the cerebellum accounted for 51% (35,128/69,174) of the total nuclei recovered. We detected known changes in the cellular composition of the four major cell types across development **(Fig. 3d)**. Purkinje cells comprised 97% (3,736/3,839) of the total nuclei recovered from the major cell types present in the cerebellar anlage at 9 PCW, then declined across development to comprise only 32% (371/1,145) at 20 PCW. At the same time, granule neurons comprised 1% (44/3,839) of the total nuclei recovered from the major cell types present in the cerebellar anlage at 9 PCW, then increased across development to comprise 58% (659/1,145) at 20 PCW. Despite its critical importance for neuron generation and requirement for proper cerebellar development, the overall size of the RL within the cerebellar anlage is small. The RL comprised only 1% (1,018/69,174) of the total nuclei recovered from the cerebellum across development, with 822 (81%) RL nuclei detected among 59,608 total nuclei recovered in our largest dataset **(Extended Data Fig. 6a and Supplementary Table 7)**.

### Molecular distinction between RL compartments (Fig. 4)

We recently determined that the human RL has unique cytoarchitectural features that are not shared with any other vertebrate, including the non-human primate, macaque.^11^ Specifically, the human RL begins as a simple progenitor niche, but then becomes compartmentalized into ventricular (RL^VZ^) and subventricular zones (RL^SVZ^) which persist until birth. The structural compartmentalization of neural progenitors into RL^VZ^ and RL^SVZ^ prompted us to select cells in the RL population, subcluster the cells, and examine the molecular correlates that define the RL^VZ^ and RL^SVZ^ compartments **(Fig. 4)**. To annotate the subclusters, we first examined the expression of classic RL markers. Indeed, *MKI67, PAX6*, and *LMX1A* were expressed throughout the subclusters, consistent with their known expression as RL markers. *WLS, 50×2*, and *CRYAB* expression was restricted, which enabled us to identify the RL^VZ^ **(Fig. 4b and Supplementary Table 10)**. One subcluster expressed *CA8*, consistent with Purkinje cells originating from the intermediate zone, and another expressed *LMX1B*, consistent with choroid plexus epithelium. We observed marked changes in the proportions of cells within RL compartments during development, with the proportion of cells occupying the RL^VZ^ generally decreasing across development and cells in the RL^SVZ^ increasing. Next, we identified additional genes with RL spatially-restricted expression. We selected the top RL markers defined by our spatial RNA-seq analysis **(Fig. 2f)** and examined expression at the singlecell level within the RL subclusters **(Fig. 4e)**. *OLIG3, R5P03*, and *5LFN13* were expressed throughout the RL while *WNT2B, CALCB, ATP6V1C2*, and *CALCA* were expressed in the RL^VZ^, and *DPYD* expression was enriched in the RL^SVZ^.

### Developmental trajectory of the rhombic lip lineage (Fig. 5)

The RL is a transient stem cell reservoir for all glutamatergic neuron progenitors in the developing cerebellum.^25,26^ Glutamatergic cerebellar nuclei neurons (eCN) are generated first, then granule cell progenitors (GCP) of the external granule layer that proliferate, differentiate, and migrate to become granule neurons of the internal granule layer arise, and unipolar brush cell (UBC) interneurons are generated last.^25,27^ We selected the RL, GCP, GN, and eCN/UBC populations, subclustered the cells, and ordered them according to pseudotime **(Fig. 5a-b)**. We used Monocle 3 ^28^ to investigate the trajectories that cell types navigate from their origin in the rhombic lip through transitional zones within the cerebellar anlage. We confirmed the predicted developmental trajectory, including temporal progression and expression of classic markers, with one branch of the RL trajectory giving rise to granule cell progenitors, then granule neurons, and the other branch giving rise to eCN/UBC **(Fig. 5b)**. Plotting relative gene expression in pseudotime, then coloring cells by cell type demonstrated the expected temporal expression associated with RL progenitors differentiating into eCN/UBC or GCP and GN **(Fig. 5c)**. Canonical RL gene expression *(MKI67, OTX2, LMX1A, EOMES)* declines as differentiate into eCN/UBC. MIK67 and DCC expression in GCP declines as GCP differentiate into GN concurrent with increased expression of *RELN* and *RBFOX3*. We selected the top markers for RL and EGL defined by our spatial RNA-seq analysis **(Fig. 2f)** and examined expression of these marker genes at the single-cell level within the RL trajectory **(Fig. 5c)**. Among the top 10 RL markers, *RSPO1, WNT2B, OLIG3, SLFN13, CALCB, CALCA*, and *ATP6V1C2* expression was largely confined to the RL, *RSP03* and *DPYD* expression was highest in RL and eCN/UBC. Among the top 10 EGL markers, expression was largely confined to the GCP, though the overall magnitude of expression was low.

Consistent with their RL origin, eCN and UBC express the classic RL markers *PAX6, LMX1A*, and *EOMES*.^27, 29^ We identified eCN/UBC on the basis of these markers and the absence of *MIK67* expression since eCN and UBC are non-proliferative at the ages sampled **(Fig. 3a-b, Supplementary Tables 8 and 9)**. The cells within this cluster originated from all ages sampled (9-21 PCW) and were distinct from other glutamatergic neurons (GCP and GN) that similarly originate from neural progenitors of the RL **(Fig. 4)**. To maximize our ability to detect differences in developmental origin between eCN and UBC, we selected cells in the eCN/UBC population from 11 PCW or 18-21 PCW, subcustered the cells, and examined the molecular correlates that define eCN and UBC **(Fig. 5)**. *PAX6, LMX1A*, and *EOMES* were expressed throughout this cluster. We next attempted to distinguish eCN and UBC by examining LMX1A/EOMES co-expression in the eDCN/UBC cluster, but only a few co-expressing cells were detectable **(Extended Data Fig. 8)**, limiting this analysis.

**Fig. 8.**
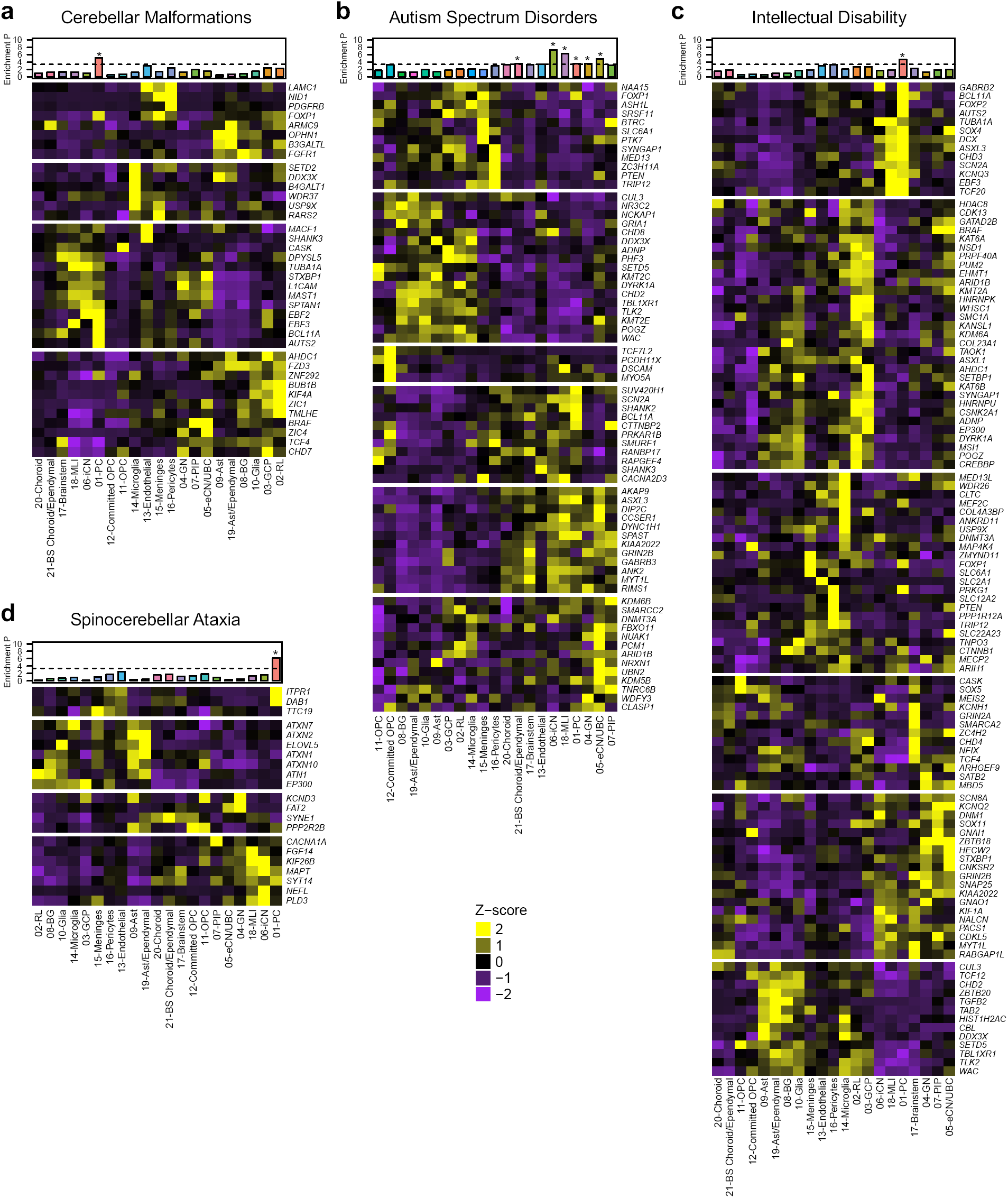
Cerebellar cell type enrichment in pediatric and adult diseases. **a-f**, Heatmaps of mean expression per fetal cerebellar cell type for genes associated with pediatric (cerebellar malformations, autism spectrum disorders, intellectual disability) or adult (spinocerebellar ataxia) diseases. Color scheme is based on Z-score distribution. In the heatmaps, each row represents one gene and each column represents a single cell type. Gene expression was clustered by row. Horizontal white lines indicate branch divisions in row dendrograms (not shown). The full list of genes is provided in Supplementary Table 11. Enrichment P values (-Log10 P value) for each cell type are shown in the top bar plots. The dashed line is the Bonferroni significance threshold. Asterisk (*) indicates significance (P<0.05).

### Purkinje cells dominate the developing cerebellar anlage (Fig. 6)

Purkinje cells arise from the ventricular zone during early embryonic stages and dominate the cerebellar anlage by 10 PCW.^30^ The Purkinje cell cluster represented the cell type with the most nuclei recovered **(Fig. 3)**. The cells within this cluster originated from all ages sampled (9-21 PCW) and were distinct from other GABAergic neurons (inhibitory cerebellar nuclei and *PAX2+* interneuron progenitors) that similarly originate from neural progenitors of the ventricular zone. To examine early markers of human Purkinje cell subtypes, we selected all cells in the Purkinje cell cluster, subclustered the cells, and then used Monocle 3 ^28^ to order them in pseudotime **(Fig. 6)**. By labeling the cells by sample age, we detected a temporal progression among Purkinje cells **(Fig. 6a)**. Plotting relative gene expression in pseudotime, then coloring cells by sample age demonstrated little fluctuation in the expression of canonical Purkinje cell marker genes, with the exception of *CALB1* and *SKOR2* **(Fig. 6b)**. *CALB1* was expressed at higher levels in later samples, while *SKOR2* expression declined with increasing sample age, consistent with its role in early Purkinje cell differentiation.^31^ *RORA* was expressed throughout the Purkinje cell cluster, as were markers that in mouse display parasagittal banding patterns of alternating Purkinje cells,^32^ including *PLCB4* and *EBF2* **(Fig. 6c)**. Few Purkinje cells expressed more mature markers, *ALDOC*and *PCP2*.

### Deconvolution of LCM and BrainSpan (Fig. 7)

LCM is a technique that provides the opportunity to harvest subpopulations of cells from precise anatomical regions of a heterogeneous tissue sample.^33,34^ However, these enriched samples are not necessarily a pure population of cells due to contamination from adjacent cell types or tissues. Therefore, we sought to directly investigate the cell type composition of our LCM samples by using the reference gene expression profiles from our snRNA-seq dataset to dissect the LCM samples. We used CIBERSORTx,^35^ a machine learning method for inferring cell-type-specific gene expression profiles, to establish a transcriptional signature for each of the 21 cell types detected in our atlas of developing human cerebellum, then estimated the relative proportions of these cell types present in each sample of our spatial transcriptional dataset. Overall, the expected cell type had the highest relative abundance in each LCM sample **(Fig. 7a and Extended Data Fig. 9)**. Cells from the RL (02-RL) were most abundant in the LCM RL samples with a median of 52%, while they represented only 7% of the cells in the EGL samples, and were absent from PCL and bulk cerebellum samples. Granule cell progenitors (03-GCP) were the most abundant cell type present in the LCM EGL samples with a median of 49%, while they represented 6.5% of the cells in the RL samples, 2% in the PCL samples, and 6% in bulk cerebellum. Purkinje cells (01-PC) were the most abundant cell type present in the LCM PCL samples with a median of 43%, while they represented 16% of the cells in bulk cerebellum and were absent from RL and EGL samples.

Next, we estimated the cell type composition of bulk cerebellar samples from BrainSpan using transcriptional signatures from our fetal cerebellar dataset or from a published adult cerebellar dataset.^36^ Overall, Purkinje cells were more abundant in bulk cerebellar samples from the fetal period of development **(Fig. 7b and Extended Data Fig. 10)**. Deconvolution using our fetal transcriptional signatures estimated that Purkinje cells comprised a median of 23% of the BrainSpan fetal samples, compared to postnatal samples with a median of 13%. Deconvolution using the adult transcriptional signatures estimated that Purkinje cells comprised a median of 16% of the BrainSpan fetal samples, compared to postnatal samples with a median of 5%. As expected, granule neurons were more abundant in postnatal samples. However, while the fetal cerebellar dataset underestimated the abundance of granule neurons present in the postnatal samples, the adult cerebellar signature overestimated the abundance of granule neurons in the fetal cerebellar samples. We examined the overlap among genes within the fetal and adult cell type signatures **(Extended Data Fig. 11)**. Among the major neuronal cell populations, including RL, GCP, GN and PC in fetal cerebellum and granule neurons and Purkinje cells in adult cerebellum, only four genes were shared between the fetal and adult transcriptional signatures. Three genes (*RAB26, RP4-665J23*.*1, RPL19)* were expressed in fetal granule cell progenitors and adult granule neurons and one gene (*MDH1)* was expressed in fetal and adult Purkinje cells. In contrast, many genes were expressed in non-neuronal cell types of both the fetal and adult cerebellum (n = 42 microglia; 25 endothelial cells; 10 oligodendrocytes).

### Cellular convergence of disease (Fig. 8)

Cerebellar involvement is central to major neurodevelopmental and adult onset neurodegenerative disorders.^1,3^ We used our atlas of developing human cerebellar cell types to identify the cell types in which mutations causing pediatric and adult diseases act as a way to provide a framework for understanding disease **(Fig. 8 and Supplementary Table 11)**. We first examined the enrichment of genes associated with structural cerebellar malformations, namely cerebellar hypoplasia and Dandy-Walker malformation, that are commonly diagnosed prenatally.^6,23^ Genes associated with these common cerebellar malformations were significantly enriched in Purkinje cells, with prominent expression of *AUT52, BCL11A, EBF2*, and *EBF3* **(Fig. 8a)**. Next, we examined the enrichment of genes that cause Joubert syndrome, a recessive neurodevelopmental ciliopathy defined by a distinctive hindbrain malformation.^37^ None of the Joubert syndrome genes showed significant enrichment in cerebellar cell types **(Supplementary Fig. 11a)**. Then, we examined the enrichment of high-confidence risk genes for ASD defined by rare, *de novo*, putatively damaging genetic variants identified in large-scale exome and whole genome sequencing studies.^38^’^43^ There was substantial variability in gene expression across cell types with significant enrichment of gene expression in multiple cell types: brainstem, inhibitory cerebellar nuclei (iCN), molecular layer interneurons (MLI), Purkinje cells, granule neurons, and eCN/UBC **(Fig. 8b)**. ASD genes were most prominently expressed in Purkinje cells *(5HANK2, SUV420H1, BCL11A, ASXL3)* and eCN/UBC (*CCSER1, NRXN1, DIP2C, FXBOll, NUAK1, UBN2, PCM1)*. We extended this analysis to examine high-confidence ID genes^42,44,45^ and found prominent expression with significant enrichment in Purkinje cells (*AUTS2, PRKG1, FOXP2, CHD3, GABRB2, BCL11A, EBF3, SOX4, TCF20, DCX, ASXL3)* **(Fig. 8c)**. Lastly, we examined the expression of genes associated with two adult-onset neurodegenerative disorders. The spinocerebellar ataxias (SCA) are progressive disorders with autosomal dominant inheritance that lead to irreversible Purkinje cell loss.^8^ SCA genes were significantly enriched in Purkinje cells, driven by *DAB1* and *ITPR1* expression **(Fig. 8d)**. Interestingly, SCA genes were prominently expressed in inhibitory cerebellar neurons (*KIF26B, MAPT, PLD3, NEFL*), consistent with neuronal atrophy of the pons and deep cerebellar nuclei identified at autopsy in SCA patients^46^ Alzheimer’s disease is a progressive disease associated with age-related cognitive decline and aberrant neuron-glial interactions.^47^ We examined the enrichment of risk genes for Alzheimer’s disease identified in a recent case-control exome sequencing study.^48^ None of the Alzheimer’s disease genes showed significant enrichment in cerebellar cell types **(Supplementary Fig. lib)**. Cell type enrichment exceeded Bonferroni significance using a down sampled dataset of 100 cells per cell type.

## DISCUSSION

Our combined approach of using spatially isolated and single-cell capture methods provide complementary transcriptional datasets that inform and validate the molecular composition of multiple cell types in the human cerebellum. Here, we provide a map of expression profiles for the major cell types present in the human cerebellum from 9 to 22 PCW. This *‘Developmental Cell Atlas of the Human Cerebellum’* can provide molecular context for comparative evolution, benchmarking *ex vivo* model systems, and investigating disease cell type origins.

Human cerebellar development is comparatively protracted, extending from 30 days post conception through the second postnatal year of life.^5,49^ The 17 week window of cerebellar development profiled here represents only a small slice of human cerebellar development. Yet this window covers crucial developmental epochs with the stereotypical lamination of the cerebellum emerging during this time.^5, 11^ We recovered all major cell types present in the developing cerebellum and even captured some rare, transient populations, such as the RL. The emerging lamination provided an anatomical scaffold that facilitated our use of LCM to specifically capture RL and EGL progenitors and differentiating cells in the PCL.

The human RL is composed of an inner RL^VZ^ and an outer RL^SVZ^ which is a feature of cerebellar development that may be unique to humans and responsible for the evolutionary expansion of the human cerebellum.^11^ Using LCM, we precisely captured cells occupying the RL by using cytoarchitechural boundaries to select the inner RL^VZ^ and outer RL^SVZ^ then molecularly profiling these subregions across mid-gestation. Both subcluster and trajectory analyses of the cell types resident in this limited RL population clearly demonstrate the ability of the single-cell data to recapitulate morphological findings, readily distinguishing the RL^VZ^ and outer RL^SVZ^ subcompartments. The RL subcluster analysis further enabled us to detect contamination of one sample by adjacent tissue. We detected choroid plexus epithelium (CPe) in one sample, likely reflecting incomplete dissection of the choroid plexus from this specimen.

During mid-gestation the RL generates cells that migrate to form the EGL and also produces UBCs, which are excitatory glutamatergic interneurons.^4^ Additional excitatory cerebellar interneurons (eCN) are also generated from the RL, but their formation concludes prior to 8 PCW.^11^ Proliferative human RL^SVZ^ progenitors express both *LMX1A* and EOMES.^11^ We also recovered a population of cells that expresses *LMX1A* and *EOMES*, but not the proliferative marker, *MKI67*, leading us to identify this cluster as eCN/UBC. While eCN express *LMX1A* but not *EOMES*, UBC express both *LMX1A* and *EOMES*. Attempts to identify colocalization of these markers and discriminate eCN and UBC resulted in too few cells (6 total). Additional experiments are required to distinguish among all of these closely related cell types, which is important because they have been implicated as the origin for group IV medulloblastoma, a poorly understood and aggressive subtype of childhood cerebellar tumor.^50,51^

Although neurogenesis in the ventricular zone concludes between 8 and 10 PCW, during early and midgestation development there is extensive migration of all ventricular zone derivatives, including Purkinje cells and PAX2+ interneuron progenitors (PIPs) through the cerebellar anlage as cerebellar morphogenesis and cerebellar circuitry are being established.^30^ In our dataset, early differentiating inhibitory interneurons (iCN) are readily distinguished from RL-derived eCN/UBCs, as are differentiating inhibitory interneurons derived from PIPs. Although Purkinje cell morphology and circuitry are nearly identical across the cerebellum, up to 50 molecularly heterogeneous Purkinje cell clusters, in part related to Purkinje cell birthdate, are present in the mouse cerebellum at late gestational stages.^52^ These Purkinje cell clusters are subsequently transformed into longitudinal stripes along the mediolateral axis and correlate with function.^32,53^ Although Purkinje cells comprised the largest cell type recovered in our dataset, we did not readily detect Purkinje cell clusters with distinct transcriptional profiles. Canonical mature Purkinje cell markers known for their expression in parasagittal stripes across the mature cerebellum (*ALDOC, PCP2)* are not yet robustly expressed in the immature Purkinje cells captured in our data **(Fig. 6c)**. This is likely because Purkinje cell maturation begins during late gestation and peaks only after birth once all GCPs in the EGL have differentiated and migrated inward to establish the IGL. Indeed, human Purkinje cells around 20 PCW display a nascent dendritic arbor, which expands considerably during late gestation and continues after birth.^30,49^

Thus far, few prior studies have characterized the transcriptional landscape of the developing human cerebellum, with only bulk RNA-seq data available from a limited number of fetal cerebellar samples within the BrainSpan resource.^16,17^ Our data significantly augment prior cerebellar data by providing 3fold more transcriptional data with spatially resolved bulk transcriptional profiles and further adding *70,000* single-nucleus transcriptomes. We provide a direct comparison of our spatial RNA-seq dataset to a bulk cerebellum RNA-seq dataset showing that there is little correspondence between our spatially resolved data and bulk cerebellar data. Only one co-expression module from BrainSpan cerebellum correlated with our fetal EGL data. We inferred cell type composition of our LCM and BrainSpan bulk cerebellum RNA-seq datasets by applying transcriptional signatures for each cell type present in our *‘Developmental Cell Atlas of the Human Cerebellum’*. Our spatially captured data showed an abundance of the targeted cell type at 40%, with abundances for other cell types at <20% **(Extended Data Fig. 9)**. When we applied these fetal cell type signatures to the cerebellum data in BrainSpan, the abundance of Purkinje cells was inferred to be about 10-20% across development, while granule neurons were estimated to comprise <10% of the cerebellum. We also applied transcriptional signatures for adult cerebellar cell types^36^ to the bulk cerebellum data in BrainSpan since the majority of these samples are derived from postnatal ages (22 out of 35 total cerebellar samples). This analysis estimated that granule neurons comprised >60% of the BrainSpan samples at all developmental stages. To reconcile these differences, we compared the fetal and adult granule cell signatures, identifying few shared genes **(Supplementary Fig. 12)**. This is not surprising since fetal granule neurons are only just beginning to emerge at the time points included in our prenatal cerebellar dataset, while the adult cerebellar dataset profiled fully mature granule cells from 35, 48, and 49 year old individuals. Thus, while bulk cerebellar data emphasize the progression of predominantly Purkinje cells prenatally to predominantly granule neurons postnatally, our *‘Developmental Cell Atlas of the Human Cerebellum’* provides molecular details of multiple cell types that are important for understanding diseases rooted in early cerebellar development.

Finally, we found that genes associated with neurodevelopmental and adult onset neurodegenerative disorders primarily implicate Purkinje cells, consistent with prior research.^1,54^ Additional data are required to close the gap from 22 PCW to 35 years and complete the cellular and transcriptional characterization of the human cerebellum across the complete human lifespan.

## MATERIALS AND METHODS

### Cerebellum samples

Specimens from fetal (9-22 PCW) human cerebellum were obtained from the Birth Defects Research Laboratory at the University of Washington or the Human Developmental Biology Resource^55^ with ethics board approval and maternal written consent. This study was performed in accordance with ethical and legal guidelines of the Seattle Children’s Hospital institutional review board.

### Histology, immunohistochemistry, and *in situ* hybridization analyses

Fixation, tissue processing, and immunohistochemistry were performed as previously described^49^ using the following primary antibodies: Calbindin (Swant, CD38, rabbit, 1:3000), PAX6 (Biolegend, 901301, rabbit, 1:300), SKOR2 (Novus, NBP2-14565, rabbit, 1:100), NEUN (Millipore, MAB377, mouse, 1:100), and TBR2 (eBioScience, 25-4877-42, mouse, 1:250). All sections were counterstained using Vectashield DAPI (H1000, Vector labs) which marks all nuclei.

*In situ* hybridization was performed using commercially available probes from Advanced Cell Diagnostics. Manufacturer recommended protocols available on the ACD webpage were used without modification. Probes used include: *LMX1A* (#540661), *MKI67* (#591771), *ATOH1* (#417861), *OTX2* (#484581), and *HOXB3* (custom-made). All sections were counterstained using Fast Green.

Slides processed for fluorescent IHC were imaged using Zeiss LSM Meta confocal microscope and ZEN 2009 software (Zeiss). Brightfield imaging was performed using a Nanozoomer Digital Pathology slide scanner (Hamamatsu; Bridgewater, New Jersey). Barring minor adjustments limited to contrast and brightness to the entire image, no additional image alteration was performed.

### Laser capture microdissection

Whole cerebellum was dissected from 16 fetal specimens that had intact calvaria to ensure correct orientation of the cerebellum. Intact cerebella were embedded in OCT, frozen at −80°C, and cryosectioned at 16-μm in the sagittal plane through the cerebellar vermis onto PEN Membrane Glass Slides (Applied Biosystems, USA). Total RNA was isolated from one whole section using the Qiagen RNeasy Micro Kit and RNA quality was assessed using the Agilent Bioanalyzer 6000 Pico Kit before proceeding with LCM. LCM was performed using the Leica DM LMD-6000 Laser Microdissection system to capture tissue containing PCL and EGL from each of 6-8 sections per slide into separate collection tubes. Total RNA was then isolated from LCM-enriched samples pooled across 9 slides using the Qiagen RNeasy Micro Kit. LCM was previously performed to capture RL ventricular (RL^VZ^) and subventricular zones (RL^SVZ^), then total RNA was isolated from RL^VZ^ and RL^SVZ^ resulting in two RNA samples per ii specimen.

### RNA-seq and analysis

Sequencing libraries were prepared using the Illumina TruSeq RNA Access Prep Kit (Illumina, USA) and 25 ng of total RNA per sample, according to the manufacturer’s protocol. RNA libraries were barcoded and sequenced including 6-8 samples per lane on an Illumina HiSeq 2000. FASTQfiles for RL^VZ^ and RL^SVZ^ samples from the same specimen (phs001908.vl.pl) were merged to generate the RL dataset and analyzed together with data for the other samples. Paired-end reads (100 bp) were aligned to the *Human* reference genome (NCBI build 37/hg19) using STAR,^56^ genes counts were summarized using HTSeq,^57^ and gene-level differential expression was analyzed using DESeq2^58^ specifying ∼ region + sex as the experimental design. Sample sex was confirmed by comparing expression of the female-specific non-coding RNA*XIST*and the chromosome Y specific gene *DDX3Y*. Significant results are reported as Benjamini-Hochberg adjusted *P* values.

### Gene co-expression network analyses

Weighted gene co-expression network analysis (WGCNA) was performed using the R package.^20^ Summarized gene counts were converted to RPKM using RNA-SeQC^59^ v1.1.8. Log2-transformed RPKM values were used for this analysis as described previously.^16^

### BrainSpan RNA-seq data

Gene-level expression data in counts and RPKM for the BrainSpan RNA-seq dataset generated from post-mortem human brain were downloaded (www.development.psychencode.org). We restricted our analysis to the cerebellum, selecting data from 35 individuals and including 3 brain regions: CBC (cerebellar cortex), CB (cerebellum), and URL (upper rhombic lip). Statistical analysis was performed using R (http://r-proiect.org/).

### SPLiT-seq method

Specimens were flash frozen in liquid nitrogen and stored at −80°C until use. Frozen tissue samples were pulverized on dry ice using a ceramic mortar and pestle. Pulverized samples were transferred to chilled microcentrifuge tubes and stored at −80°C until use. Nuclei were isolated from either 150 mg of pulverized tissue or an entire amount of dissected cerebellum using a published protocol.^60^ Nuclei were fixed according to the SPLiT-seq protocol.^19^ The SPLiT-seq method was performed in an initial experiment as previously described.^23^ Two additional SPLiT-seq experiments were performed using nuclei isolated from 13 cerebellar specimens. A detailed experimental protocol can be found here: https://sites.google.com/uw.edu/splitsea. Libraries were first sequenced on an Illumina NextSeq using 150 nucleotide kits and paired-end sequencing. Libraries were then sequenced on an Illumina NovaSeq S2 flow cell by SeqMatic (experiment 2) or the Northwest Genomics Center at the University of Washington (experiment 3). We used the SPLiT-seq pipeline to convert FASTQ files into digital gene expression matrices from each sequencing run: https://github.com/vizhang/split-seq-pipeline.

### snRNA-seq analysis

Deep and shallow sequencing runs from experiment ^l,23^ and shallow sequencing runs from experiment 2 and experiment 3 were filtered independently (datasets 1K, 5K, 10K, and 80K, respectively). Cells with <200 genes, >4 standard deviations above the median number of genes or UMIs, or >1-5% mitochondrial genes were removed from the analysis **(Supplementary Table 7)**. DoubletFinder^22^ was used to detect likely doublets assuming a rate of 5%, that were discarded from analysis. Sample sex was confirmed by counting reads mapped to the female-specific non-coding RNA *XIST* and the chromosome Y specific gene *DDX3Y*. We used Seurat^24^ v3 for downstream analysis. The four filtered datasets (1K, 5K, 10K, and 80K) were combined into a single dataset using canonical correlation analysis with anchors (‘FindIntegrationAnchors’) to correct for batch effects.^61^ The top 2,000 most variable genes were used to find individual cells in each sequencing run that originate from the same biological state, which became the anchors to merge runs together. The resulting dataset was then scaled and centered as well as regressed out cell cycle difference (S.score – G2M.score). Data dimensionality of the integrated dataset was reduced by principal component analysis (‘RunPCA’), then uniform manifold approximation and projection (‘RunUMAP’), then Shared Nearest Neighbor (SNN) Graph construction (‘FindNeighbors’), and finally Louvain clustering (‘FindClusters’) using the first 75 principal components and a resolution of 1.5 to determine cluster assignment. A Wilcoxon rank sum test was performed to identify differentially expressed genes for each cluster (‘FindAllMarkers’) and compare them to known gene markers for cell type assignment. One cluster with no significant differentially expressed genes and another cluster with an enrichment of mitochondrial genes were removed.

Subcluster analysis was performed by subsetting populations of interest from the overall dataset. Clustering and differential gene tests were repeated with a subpopulation specific number of principal components determined by ‘ElbowPlot’. Then, pseudotime analysis was performed using Monocle 3.^28^ Subsets were normalized (‘preprocess_cds’), dimension reduction was applied (‘reduce_dimension’), followed by clustering (‘cluster_cells’) and visualization (‘learn_graph’). Pseudotemporal ordering of cells was performed (‘order_cells’) by selecting a biologically relevant starting point. Genes of interest were used to construct the pseudotime trajectory (‘plot_genes_in_pseudotime’).

### Cell type deconvolution

We used CIBERSORTx^35^ to estimate the cell type composition in the LCM-isolated and BrainSpan RNA-seq samples. We down sampled our integrated snRNA-seq dataset to 100 cells per cell type, built a cell type signature matrix with this digital expression matrix, and imputed cell fractions for each of the 57 LCM RNA-seq samples and 35 BrainSpan cerebellar samples. We also downloaded the count matrix for a published adult cerebellar snRNA-seq dataset,^36^ from the Gene Expression Omnibus (GSE97942). We down sampled the adult dataset to 100 cells per cell type, then used this digital expression matrix to build cell type signatures for the adult cerebellum. We then applied the adult cerebellar cell type signatures to infer cell fractions in bulk cerebellar data from BrainSpan.

### Gene set curation

Disease gene lists are provided in Supplementary Table 11. The cerebellar malformation gene list was obtained from exome sequencing analysis and published Dandy-Walker malformation and cerebellar hypoplasia genes.^6,23,62^ The CBLM list included 54 genes. The Joubert syndrome gene list was compiled from published Joubert syndrome genes.^37^ The JS list included 39 genes. The ASD gene set was compiled by selecting high-confidence ASD genes identified through exome and genome sequencing.^38^’^43^ The final ASD list included 108 genes. The intellectual disability (ID) gene list was compiled by selecting genes identified through exome sequencing.^42,44,45^ The final ID list included 186 genes. The spinocerebellar ataxia (SCA) gene set was compiled by selecting genes from OMIM phenotype PS164400. The SCA list included 44 genes. The Alzheimer’s disease gene list was compiled by selecting genes identified through exome sequencing.^48^ The ALZ list included 120 genes.

### Cell type enrichment analysis

We used a one sample Z-test^63^ to identify cell types that showed enriched gene expression associated with particular gene sets. We calculated the average expression for each gene per cell type, then removed genes with expression values < 1 for more than one cell type to define a population size of 4,258 genes. Enrichment P values were corrected for multiple testing using the Bonferroni method calculated over all cell types and gene lists used [P < 0.05/(21 x 6)].

## Data and materials availability

Human tissue used in this study was provided by the Joint MRC/Wellcome (MR/R006237/1) Human Developmental Biology Resource (HDBR), UK (www.hdbr.org) and the Birth Defects Research Laboratory (BDRL), USA and covered by material transfer agreements between SCRI and HDBR/BDRL. Samples may be requested directly from the HDBR/BDRL. Processed data are available through the Human Cell Atlas (https://www.covid19cellatlas.org/aldinger20). Sequence data were deposited into the Database of Genotypes and Phenotypes (dbGaP), under accession numbers phs001908.vl.pl and XXX [in process].

## Acknowledgements

This study was funded by the National Institutes of Health under National Institute of Neurological Disorders and Stroke (NINDS), National Institute of Child Health and Human Development (NICHD), and National Institute of Mental Health (NIMH) grant numbers NS095733 to K.J.M., HD000836 to I.A.G., NS050375 to W.B.D, and MH110926, MH116488, and MH106934 to N.S. The project that gave rise to these results received the support of a fellowship from “la Caixa” Foundation (ID 100010434) to G.Sa. The fellowship code is LCF/BQ/PH9/11690010.

## Author contributions

K.A.A conceived the project, designed experiments, analyzed data, and wrote the manuscript. Z.T. performed experiments and analyzed data. P.H. performed experiments, contributed to data interpretation and manuscript preparation. M.D., M.H., L.M.O. performed experiments. M.H., C.R., A.B.R, and G.Se. provided SPLiT-seq expertise and experimental support. A.E.T., G.Sa., and B.L.G. analyzed data. F.O.G., D.O’D., and P.A. provided experimental and/or analysis support. S.N.L, N.S., W.B.D, D.D, and I.A.G supervised experiments and/or data analysis. K.J.M. provided general oversight, data interpretation, and manuscript preparation.

