## Supplemental Figures 1-12 for "Spatial and single-cell transcriptional landscape of human cerebellar development"

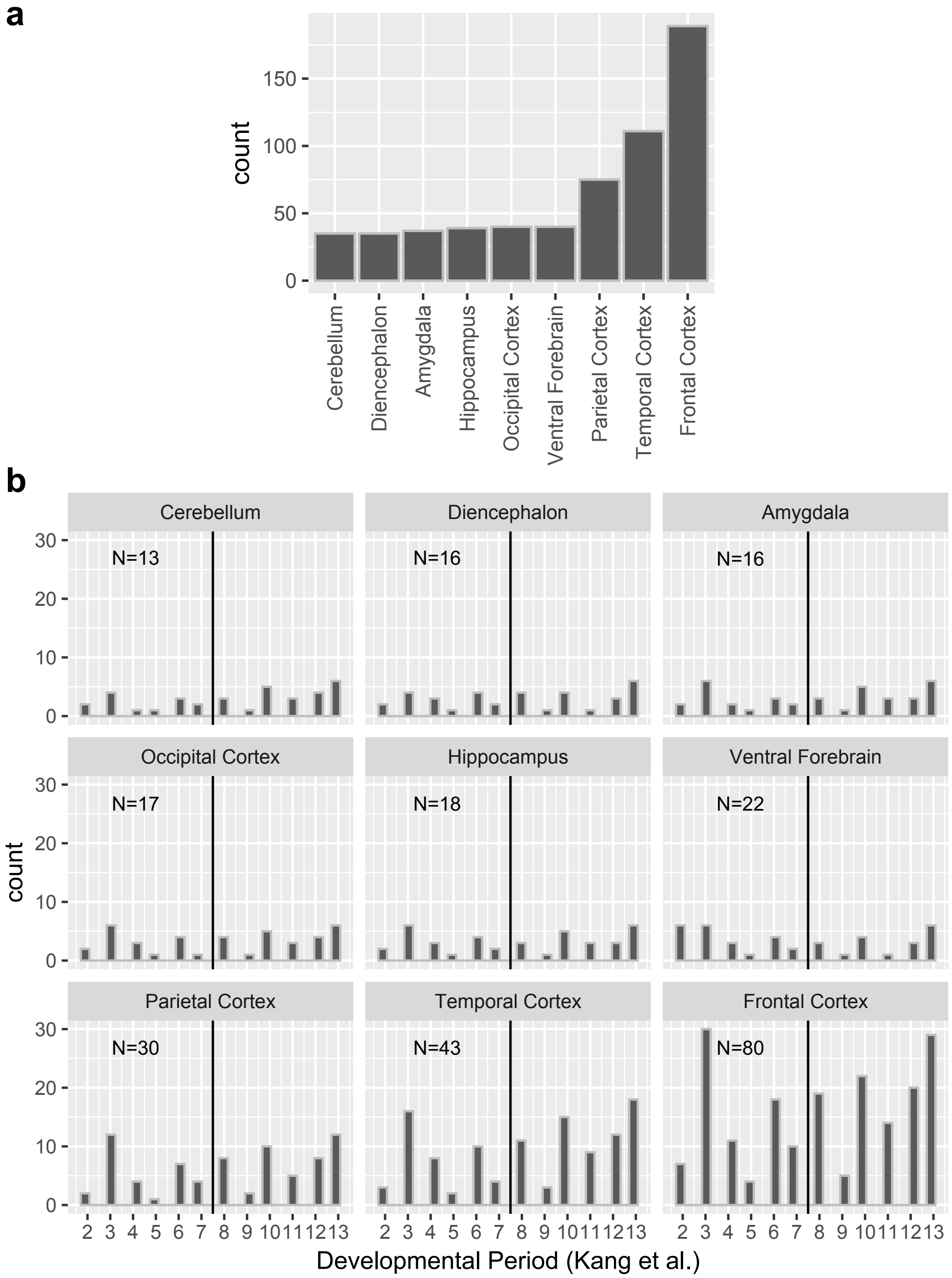


**Extended Data Fig. 1 | BrainSpan RNA-seq samples by brain region. a,** Histogram of sample counts per brain region included in the BrainSpan Atlas of the Developing Human Brain RNA-seq dataset (Li *et al.*, 2018). Brain regions are in ascending order by sample number. **b,** Histograms of sample counts by 13 developmental periods as defined by Kang *et al.*, 2011 for each brain region. Vertical black line separates prenatal (1-7) and postnatal developmental periods (8-13). Total sample count for each region during prenatal development is shown (N).

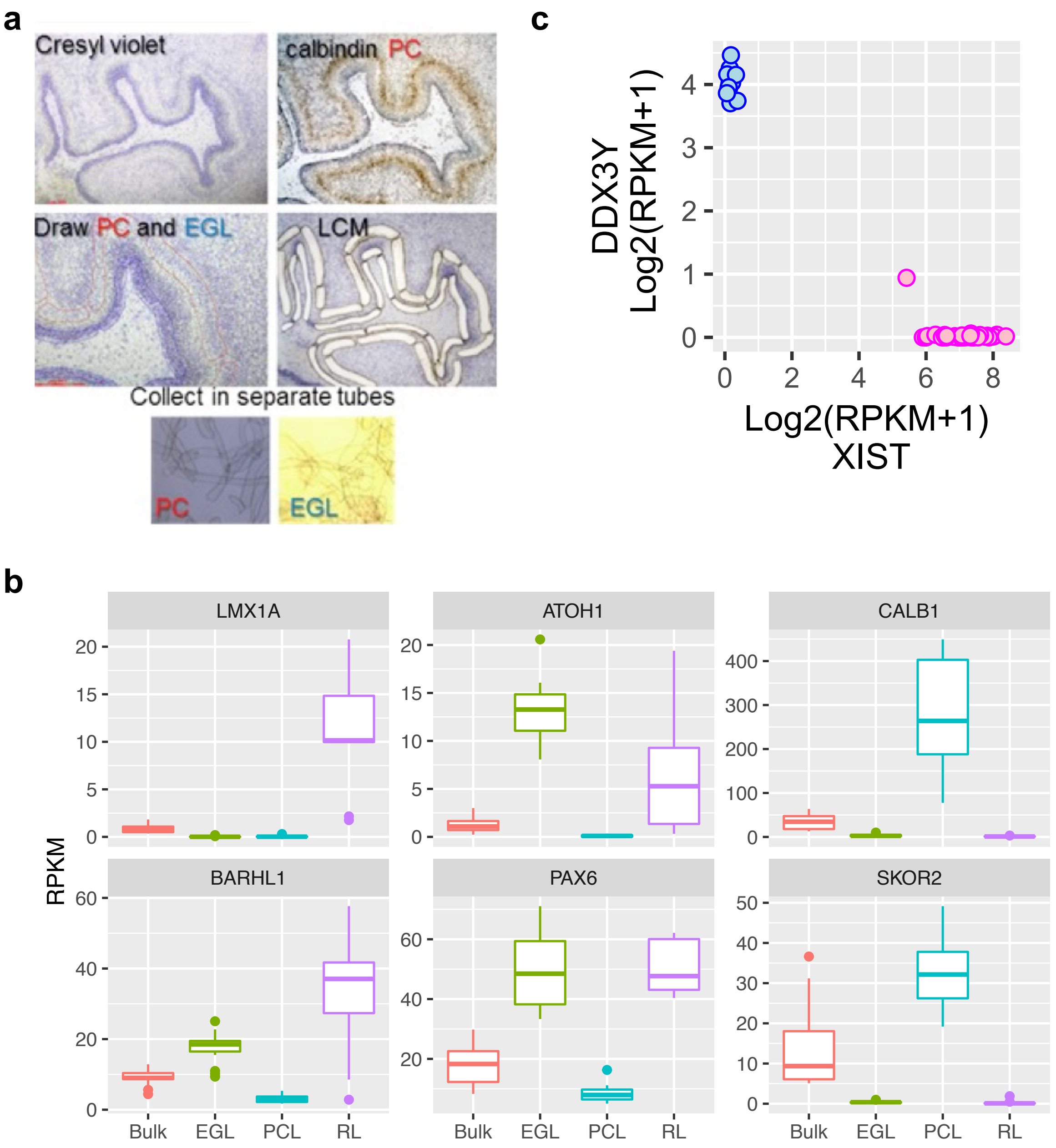


**Extended Data Fig. 2 | Quality control related analyses of LCM RNA-seq data. a,** Example of cerebellum section stained with cresyl violet and anti-calbindin antibody. Section before and after LCM, and images of tissue captured into collection tubes are shown. **b,** Boxplots of gene expression for established markers showing highest expression in the expected samples. **c,** Expression of the female-specific non-coding RNA *XIST* and the chromosome Y specific gene *DDX3Y* show correct sex assignment for female (pink) and male (blue) samples.

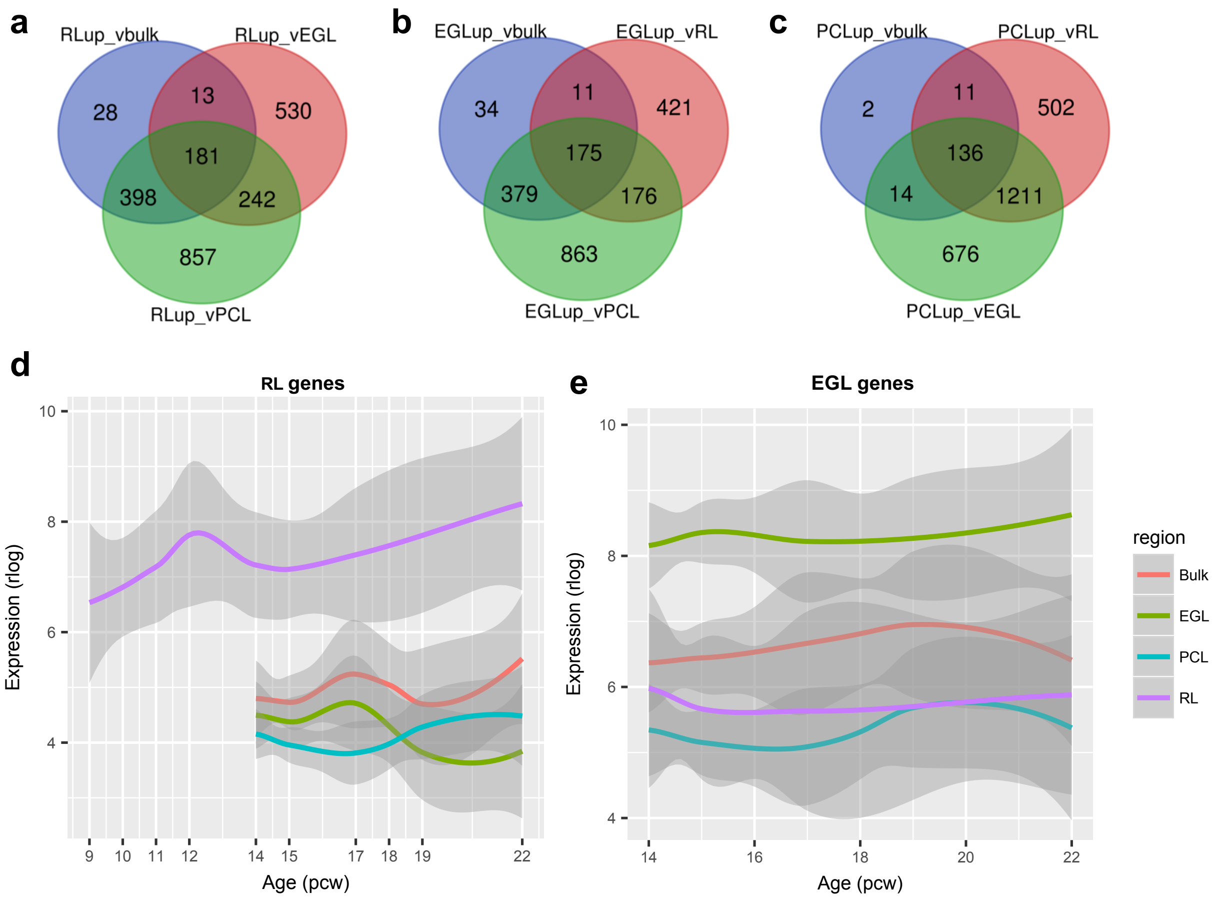


**Extended Data Fig. 3 | Spatiotemporal gene expression in the fetal cerebellum. a-c,** Venn diagrams showing the distribution of significantly differentially expressed genes among pair-wise comparisons between LCM-isolated regions and bulk cerebellum. The number of genes with region-specific expression was determined by the overlap among differentially expressed genes (N=181 for RL; 175 for EGL; 136 for PCL). **d-e,** LOESS expression values across development are shown with 95% CIs for RL-specific (d) and EGL-specific (e) genes. Spatial regions are distinguished by colors.

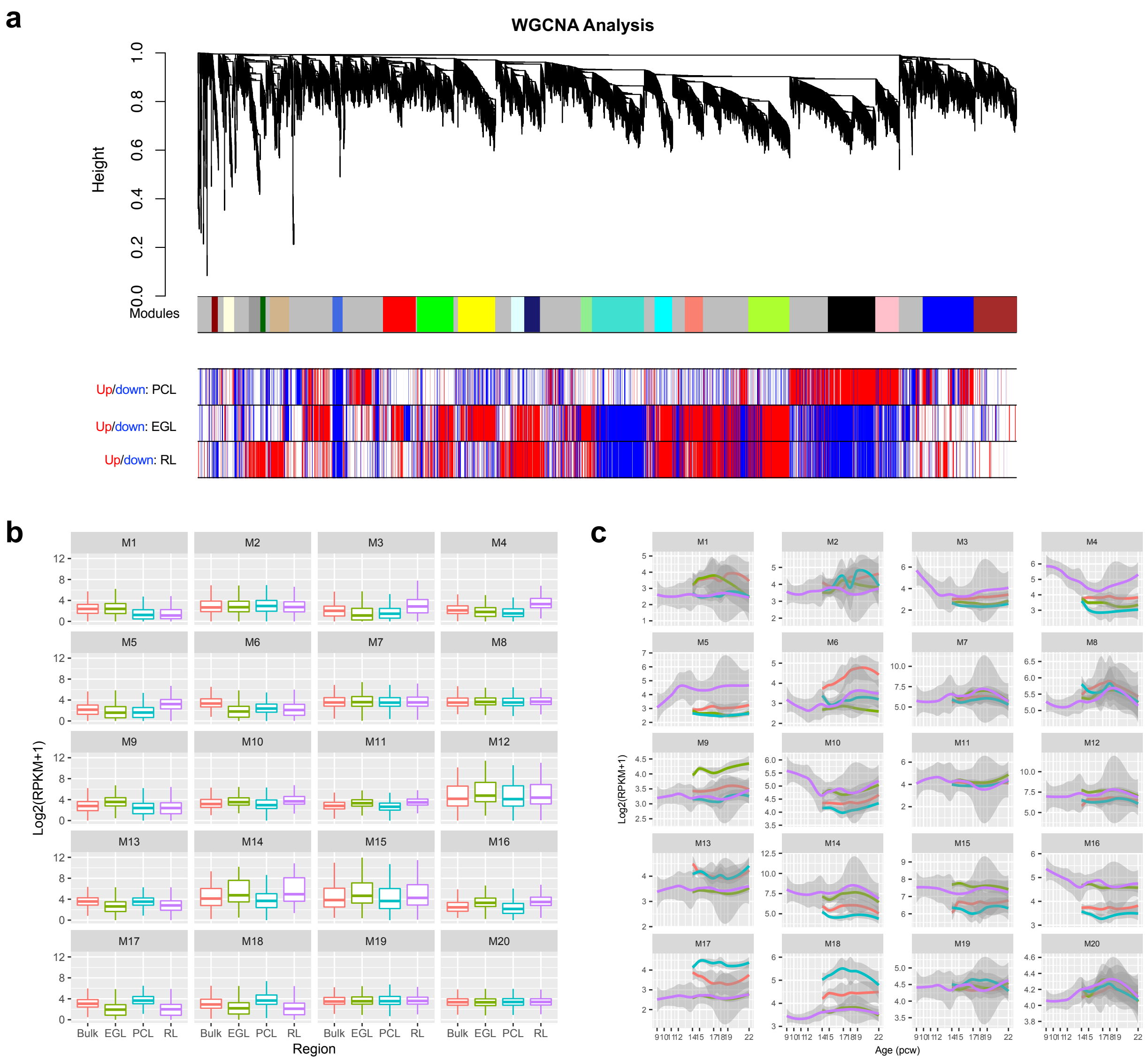


**Extended Data Fig. 4 | Co-expression modules in the developing human cerebellum. a,** Weighted gene co-expression network analysis (WGCNA) dendrogram identified 21 modules comprised of 6,336 expressed genes (row 1). M0 (grey) comprised of nonclustered genes was not analyzed further. Rows 2-4 show differential expression relationships between module genes and LCM-enriched region compared to bulk expression. **b**, Boxplots of gene expression per module for bulk and spatial regions. **c**, LOESS expression values across development are shown with 95% CIs per module. Spatial regions are distinguished by colors as in **b**. EGL, external granule cell layer; PCW, postconceptional week; PCL, Purkinje cell layer; RPKM, reads per kilobase of transcript per million mapped reads; RL, rhombic lip.

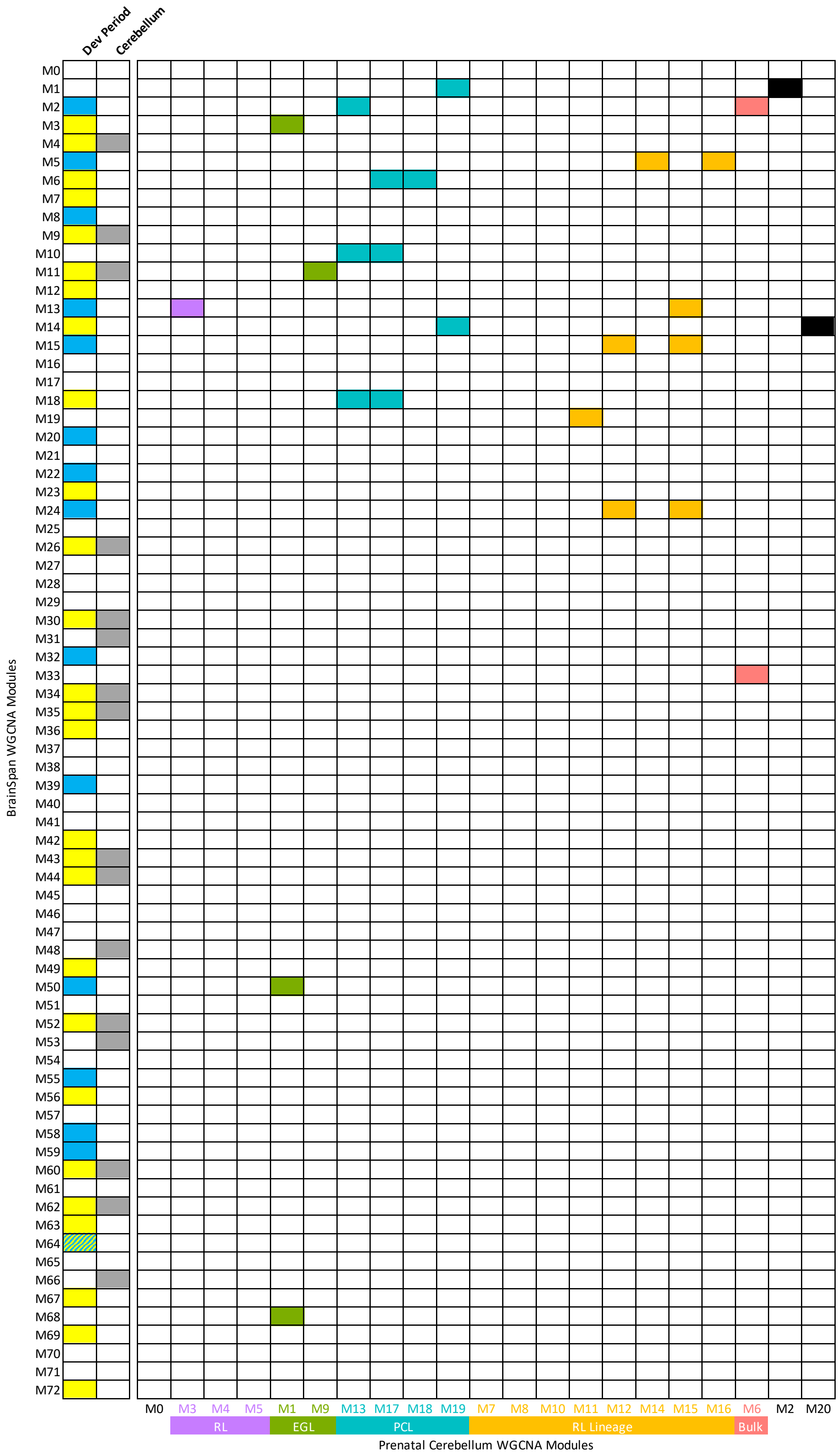
Dev Period
Cerebellum
BrainSpan WGCNA Modules
M0
M1
M2
M3
M4
M5
M6
M7
M8
M9
M10
M11
M12
M13
M14
M15
M16
M17
M18
M19
M20
M21
M22
M23
M24
M25
M26
M27
M28
M29
M30
M31
M32
M33
M34
M35
M36
M37
M38
M39
M40
M41
M42
M43
M44
M45
M46
M47
M48
M49
M50
M51
M52
M53
M54
M55
M56
M57
M58
M59
M60
M61
M62
M63
M64
M65
M66
M67
M68
M69
M70
M71
M72
M0 M3 M4 M5 M1 M9 M13 M17 M18 M19 M7 M8 M10 M11 M12 M14 M15 M16 M6 M2 M20
RL
EGL
PCL
RL Lineage
Bulk
Prenatal Cerebellum WGCNA Modules

**Extended Data Fig. 5 | Correlations between developing human cerebellum and Brainspan co-expression modules.** BrainSpan modules are represented in rows with prenatal expression yellow, postnatal expression blue, and cerebellar modules gray. Developing human cerebellum modules are represented in columns. Modules with significant enrichment (Fisher's exact test) between BrainSpan and prenatal cerebellum are shaded by color corresponding to the developing cerebellar region associated with the module. BrainSpan modules with high expression prenatally (blue), postnatally (yellow), or in cerebellum (grey) are indicated.

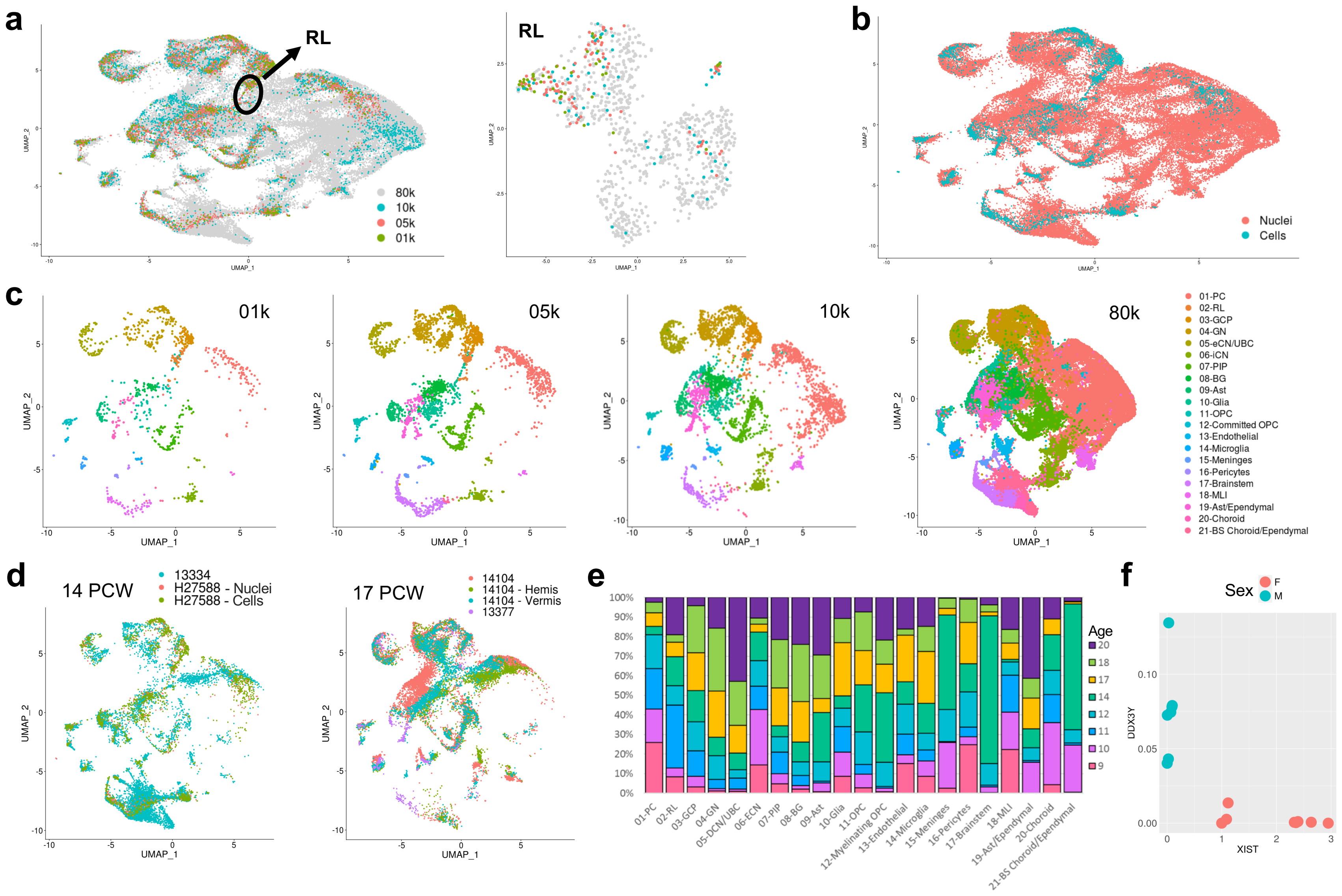

a
RL
80k
10k
05k
01k
UMAP_1
UMAP_2
b
Nuclei
Cells
c
01k
05k
10k
80k
01-PC
02-RL
03-GCP
04-GN
05-eCN/UBC
06-iCN
07-PIP
08-BG
09-Ast
10-Glia
11-OPC
12-Committed OPC
13-Endothelial
14-Microglia
15-Meninges
16-Pericytes
17-Brainstem
18-MLI
19-Ast/Ependymal
20-Choroid
21-BS Choroid/Ependymal
d
14 PCW
13334
H27588 - Nuclei
H27588 - Cells
17 PCW
14104
14104 - Hemis
14104 - Vermis
13377
e
Age
20
18
17
14
12
11
10
9
01-PC
02-RL
03-GCP
04-GN
05-DCN/UBC
06-ECN
07-PIP
08-BG
09-Ast
10-Glia
11-OPC
12-Myelinating OPC
13-Endothelial
14-Microglia
15-Meninges
16-Pericytes
17-Brainstem
18-MLI
19-Ast/Ependymal
20-Choroid
21-BS Choroid/Ependymal
f
Sex
F
M
DDX3Y
XIST

**Extended Data Fig. 6 | Quality control related analyses of snRNA-seq data. a,** UMAP visualization of 71,553 human cerebellar nuclei colored by dataset. Rhombic lip (RL) is circled. UMAP visualization of 1,018 RL nuclei colored by dataset at right (nuclei numbers: n = 41 for 1K; 88 for 5K; 67 for 10K; 822 for 80K). **b,** The same UMAP as in **a** with nuclei colored by type (nuclei or cells). **c,** The same UMAP as in **a** and **b** showing only nuclei from each dataset. Nuclei are colored by cell type. **d,** The same UMAP as in **a-c** showing only nuclei sampled from same age biological and technical replicates. **e,** Percentage of age sampled in each of the 21 cell types. Bar colors represent age sampled in postconceptional weeks (9-20 PCW). **f,** Expression of the female-specific non-coding RNA *XIST* and the chromosome Y specific gene *DDX3Y* show correct sex assignment for female (salmon) and male (turquoise) samples.

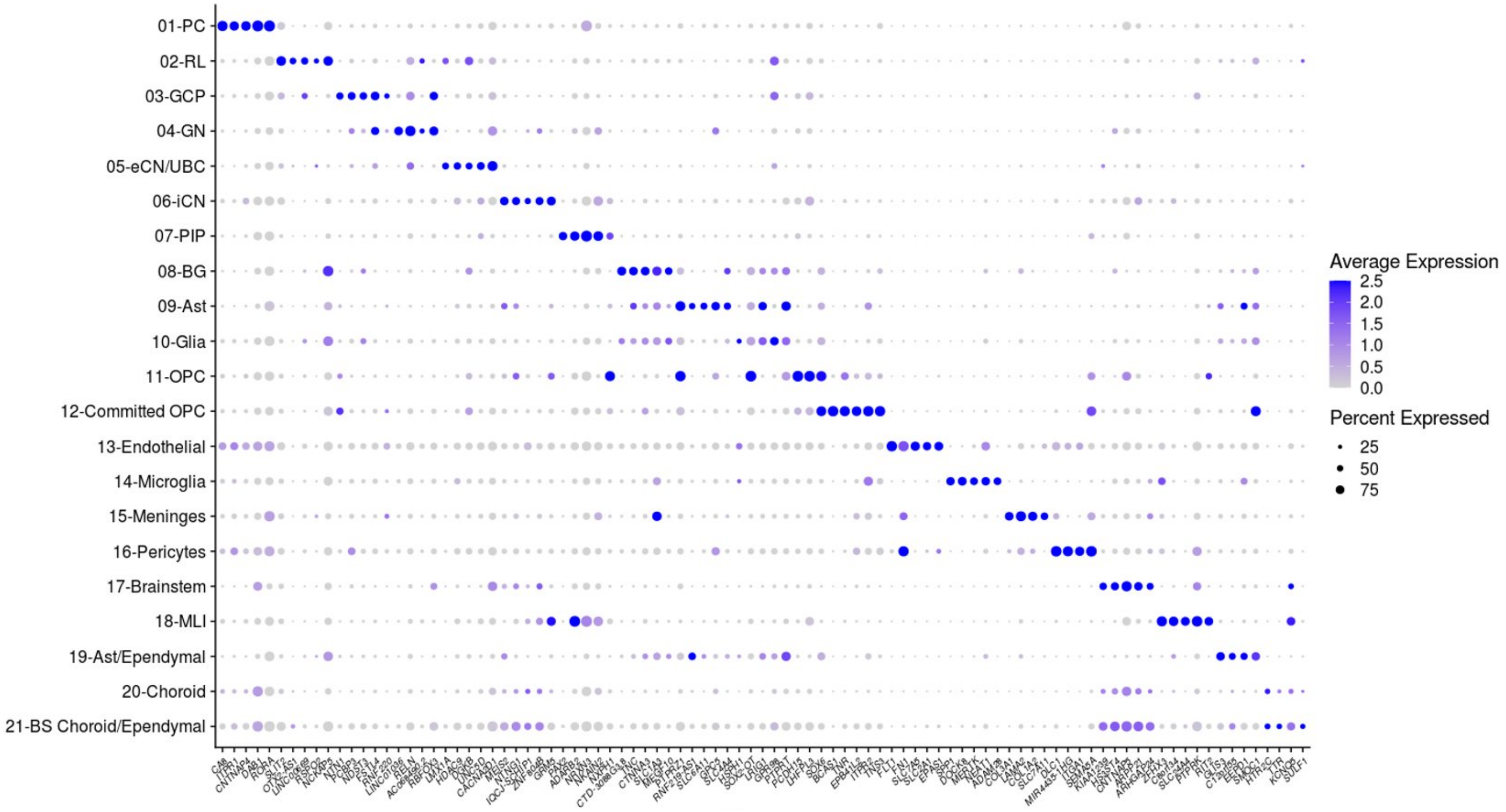


**Extended Data Fig. 7 | Cell-type-specific marker genes.** Dot plot showing expression of the top 5 most differentially expressed genes for each of the 21 cell types identified in early and mid-gestation fetal cerebellum. The size of the dot represents the percentage of cells within a cell type in which that gene was detected and its color represents the average expression level.

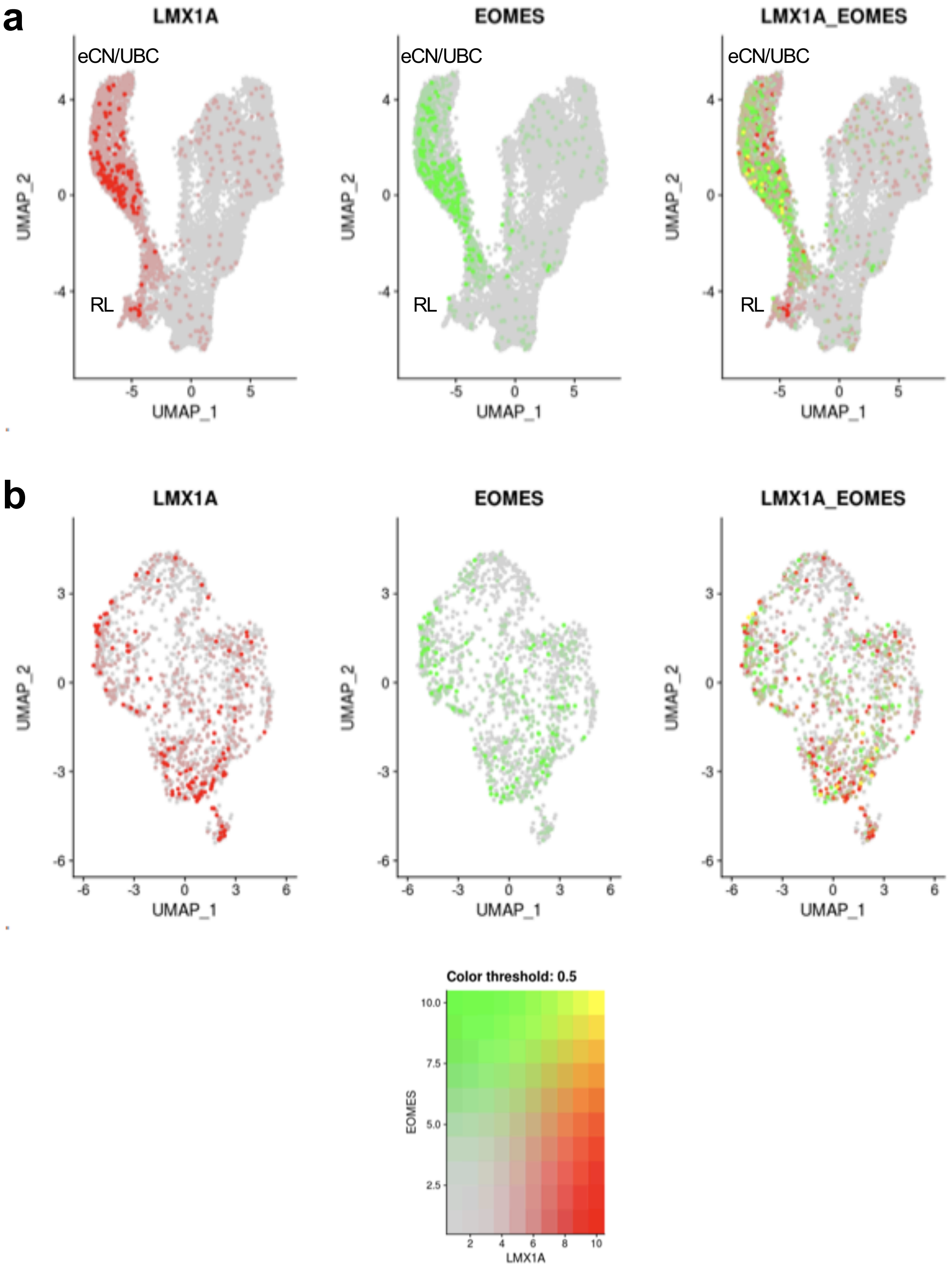
a
LMX1A
EOMES
LMX1A_EOMES
eCN/UBC
RL
UMAP_1
UMAP_2
b
LMX1A
EOMES
LMX1A_EOMES
UMAP_1
UMAP_2
Color threshold: 0.5
EOMES
LMX1A

**Extended Data Fig. 8| Co-expression of marker genes in eCN/UBC. a,** The same UMAP visualization of cell types that originate from the RL as in **Fig. 5a** with nuclei colored by expression level for *LMX1A* (red), *EOMES* (green), and co-expression (yellow). **b,** The same UMAP visualization the eCN/UBC subcluster as in **Fig. 5e** with nuclei colored by expression level for *LMX1A* (red), *EOMES* (green), and co-expression (yellow).

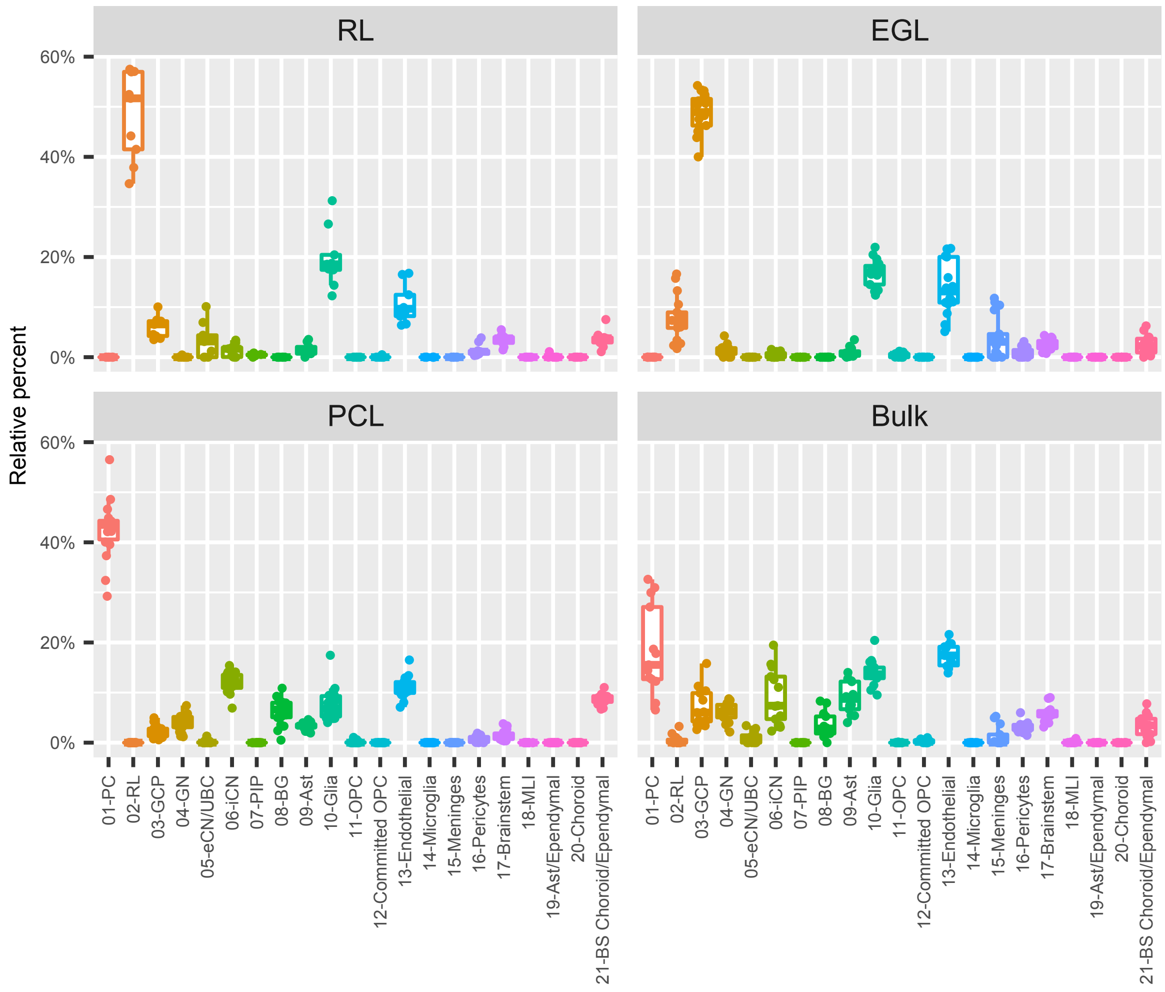


**Extended Data Fig. 9 | Cell type heterogeneity in LCM-isolated regions of the cerebellum** Box plots with data points showing the proportion of each of the 21 cell types from the *Developmental Cell Atlas of the Human Cerebellum* represented in the LCM RNA-seq data, grouped by LCM-isolated region. RL, rhombic lip; EGL, external granule cell layer; PCL, Purkinje cell layer.

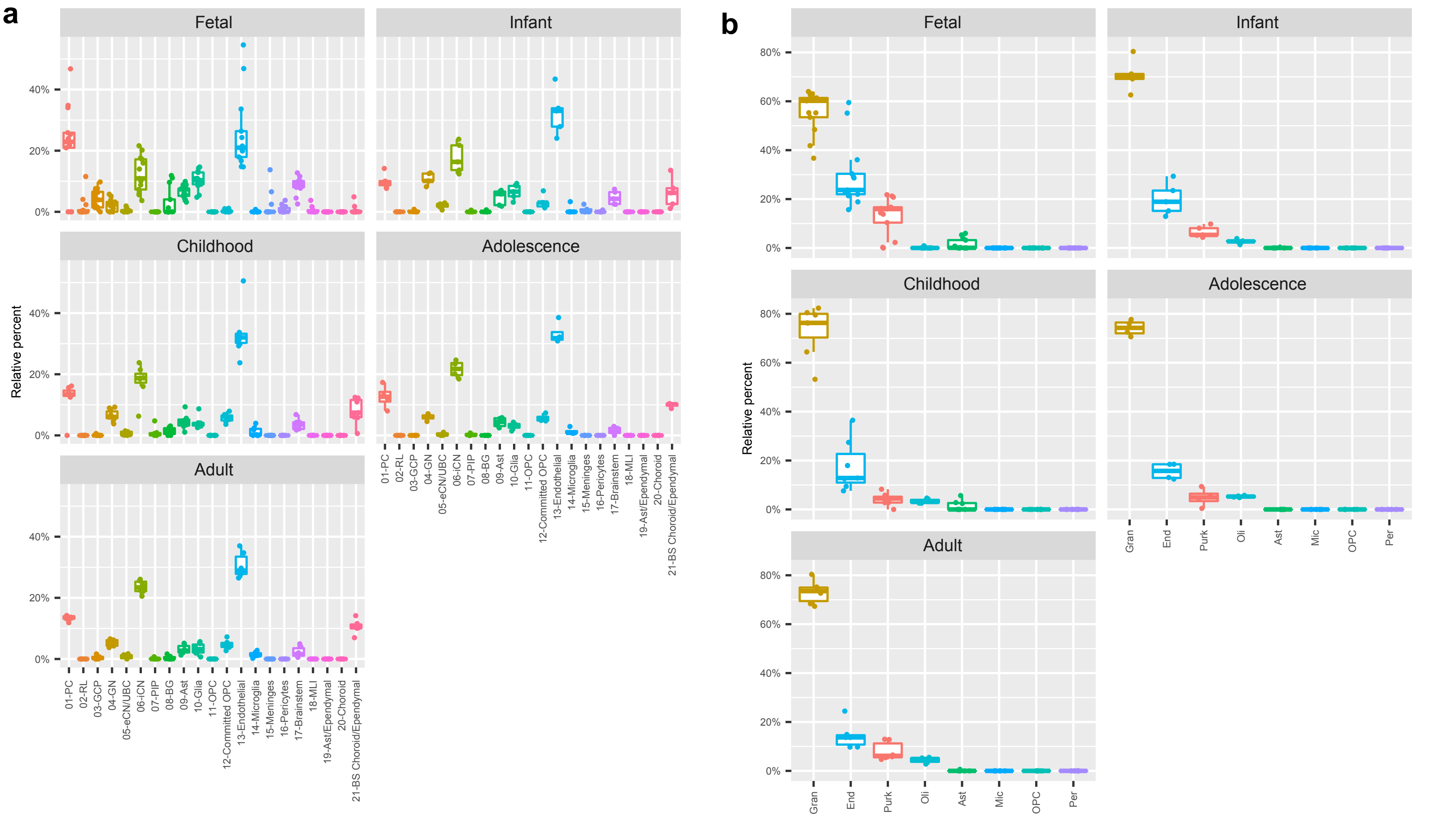


**Extended Data Fig. 10 | Cell type heterogeneity in BrainSpan cerebellum.** Box plots with data points showing the proportion of cerebellar cell types present in cerebellar samples from BrainSpan. Samples are grouped by developmental stage. **a,** Deconvolution of BrainSpan cerebellar samples using 21 cell types from *Developmental Cell Atlas of the Human Cerebellum* as the reference to estimate the relative percent of each cell type present in each BrainSpan cerebellum sample. **b,** Deconvolution of BrainSpan cerebellar samples using the 8 cell types from an adult cerebellum snRNA-seq dataset.[3]

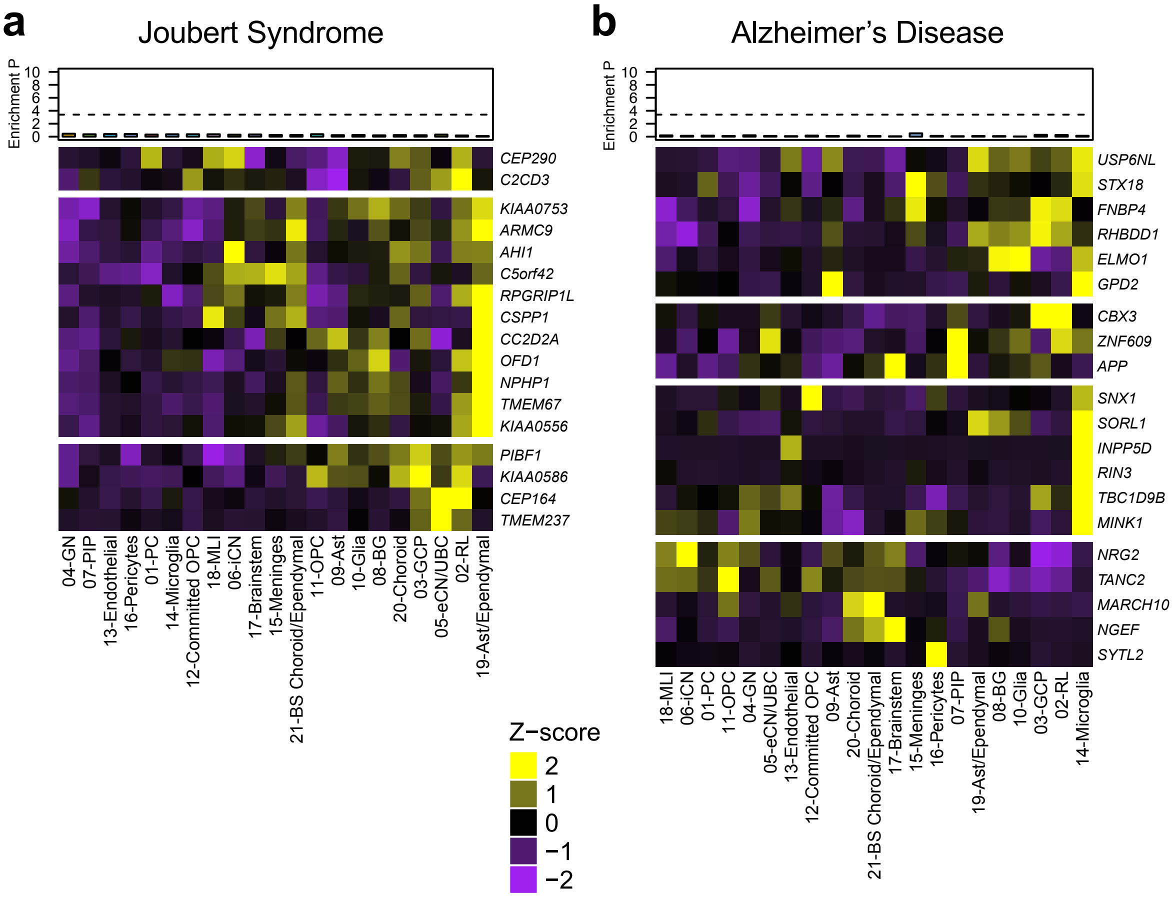


**Extended Data Fig. 11 | Cerebellar cell type enrichment in Joubert syndrome and Alzheimer's disease. a-f,** Heatmaps of mean expression per fetal cerebellar cell type for genes associated with Joubert syndrome (a) or Alzheimer's disease (b). Color scheme is based on Z-score distribution. In the heatmaps, each row represents one gene and each column represents a single cell type. Horizontal white lines indicate branch divisions in the clustering dendrograms (not shown). The full list of genes is provided in Supplementary Table 11. Enrichment P values (-Log10 P value) for each cell type are shown in the top bar plots. The dashed line is the Bonferroni significance threshold. Asterisk (*) indicates significance (P<0.05).

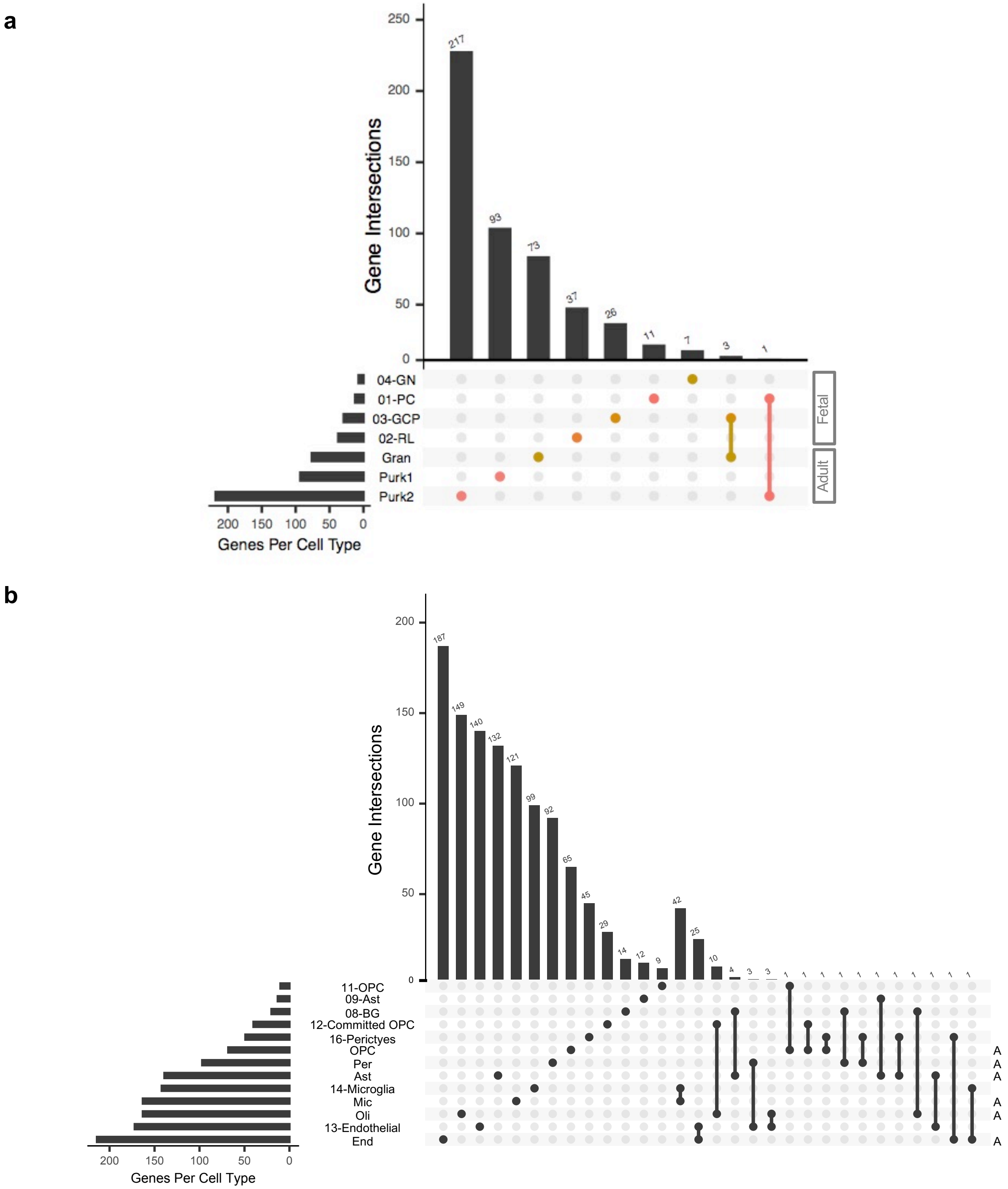


**Extended Data Fig. 12 | Intersection of genes among cell type signatures.** Cell type signature genes were selected for the indicated cell types from the *Developmental Cell Atlas of Human Cerebellum* and from adult cerebellar cell types in Lake *et al*., 2018. Bars represent the size of intersections. Dots indicate genes in each cell type. Vertical lines indicate intersections of genes between cell types. **a**, Intersection among genes expressed by major neuronal cell types in fetal and adult cerebellum. **b**, Intersection among genes expressed by non-neuronal cell types in fetal and adult cerebellum. Adult cell types are indicated (A).
